# Cryo-EM as latent structural landscape microscopy

**DOI:** 10.64898/2026.04.10.717737

**Authors:** Haizhao Dai, Qihe Chen, Lingqi Li, Yingjun Shen, Zhenyang Xu, Minzhang Li, Yufan Xie, Jingjing Zheng, Zhijie Liu, Liping Sun, Yuan Pei, Jiakai Zhang, Jingyi Yu

## Abstract

A protein populates a landscape of structural states, abundant snapshots of which are sampled in every cryo-EM experiment. Current analysis averages these snapshots into single density maps or partitions them into discrete classes, discarding the continuous dynamics encoded in the data. Continuous latent-space methods offer a promising alternative, yet whether their learned representations are physically grounded remains unresolved. Here, we realize cryo-EM as structural landscape microscopy, in which latent density directly reflects the probabilistic distribution of molecular states. A central question is whether such a landscape reflects physical reality. For integrin ***αvβ*8**, the learned landscape shows strong agreement with independently derived molecular dynamics simulations, supporting its physical plausibility. We then apply the landscape to structural states that conventional cryo-EM cannot resolve. For LIS1-mediated dynein activation, the landscape reveals a spectrum of states from dominant conformations to low-population intermediates defined by distinct binding modes, including a previously unresolved state. For the KCTD5/CUL3^NTD^/G***βγ*** complex, the landscape resolves continuous conformational pathways directly from experimental data. Probability-guided particle selection further improves reconstruction quality.

## 1 Introduction

Since the middle of the last century, modeling protein structure has relied on a single, explicit representation defined by atomic coordinates, as resolved by advancing imaging apparatus from X-ray crystallography to cryo-EM. These models have transformed our understanding of molecular function and enabled structure-guided drug design, yet they largely capture a single state of molecules that naturally adopt diverse structural forms under physiological conditions. In theory, single-particle cryo-EM readily captures this structural variability as millions of molecular snapshots. Yet, standard reconstruction approaches assume a single or a few underlying states, averaging the captured variation into a single density map that best fits the data. Discrete classification methods [1, 2] partially resolve this by partitioning single or multiple clusters of particles into a small number of states, but intermediate states and continuous dynamics remain unresolved. Recent advances in heterogeneity analysis [3–8] encode structural variability into learned latent spaces, enabling exploration of the full range of conformational variation captured in the data. However, the intrinsic structure of these latent spaces, including probability density and transition pathways, remains qualitative rather than physically grounded. Without such grounding, there is no principled basis for distinguishing genuine conformational states from noise or for identifying the transition pathways between states.

Here, we set out to realize cryo-EM as structural landscape microscopy, where the captured data directly reveals the probabilistic distribution of conformational states. The central challenge is that cryo-EM particle images have extremely low signal-to-noise ratios [9], demanding latent representations that reliably separate structural signal from imaging noise. To address this, we present CryoUNI, a universal encoder for cryo-EM particle images. Trained on large-scale diverse particle datasets, CryoUNI learns to extract robust structural representations from noisy images, mapping them into a probabilistic latent space in which density reflects the distribution of conformational states, thereby defining a relative energy landscape described by Boltzmann statistics. Within this latent space, distinct structural states emerge as density peaks, i.e., energy basins, and transitions between them follow continuous pathways. To systematically analyze this landscape, we introduce WAVE (Watershed Analysis of Variational Embeddings), which automatically identifies energy basins and their transitions from latent density, resolving both dominant conformations and rare intermediates without requiring a predefined number of states.

A central question for any learned conformational landscape is whether it reflects true physical reality. Molecular dynamics (MD) simulations offer an independent reference of this ’ground truth’, as they sample conformational space from first principles. We compared the learned landscape with long-time scale allatom MD simulations of integrin *αvβ*8, a widely used benchmark for cryo-EM heterogeneity analysis whose continuous leg rotation provides a well-defined, physically parameterizable degree of freedom. The learned latent coordinates quantitatively correspond to the angular coordinates governing this motion, establishing, to our knowledge, the first direct link between a cryo-EM latent space and an MD-derived thermodynamic landscape.

Having established the physical basis, we next turned to protein complexes whose function hinges on conformational states that conventional cryo-EM cannot resolve — transient intermediates in dynamic assemblies and continuous motions in intrinsically flexible complexes. For LIS1-mediated dynein activation [10, 11], an essential intracellular transport machinery regulated through varying complex subunit assemblies, the learned landscape uncovers low-population intermediates that are invisible to standard discrete classification yet critical to understanding the stepwise activation mechanism. For the KCTD5/CUL3^NTD^/G*βγ* complex [12–14], a ubiquitin ligase assembly whose intrinsic flexibility is averaged out under conventional reconstruction, the landscape resolves continuous conformational trajectories directly from cryo-EM data, to the best of our knowledge, for the first time. Furthermore, the physically grounded landscape enables probability-guided particle selection that substantially improves reconstruction resolution, demonstrating its utility not only for discovering hidden conformational states but also for enhancing structure determination itself.

## 2 CryoUNI learns a latent structural landscape from cryo-EM

Cryo-EM particle images carry structural information at extremely low signal-to-noise ratios, demanding representations that reliably separate structural signal from imaging noise. We address this by pretraining a universal encoder, CryoUNI, on CryoCRAB-Particle-22M (Fig. 1a), a dataset of 22 million cryo-EM particles spanning five SCOP2 structural classes [15], with molecular weight ranging from below 250 kDa to above 1 MDa. To ensure that the encoder extracts a clean structural signal rather than imaging noise, we adopt a self-supervised training objective [16, 17] customized for cryo-EM particle images. Specifically, CryoUNI is pretrained to denoise particles from one half-dataset using the complementary half as supervision, exploiting the fact that both halves share identical underlying structure but distinct noise (Fig. 1b).

**Fig. 1.**
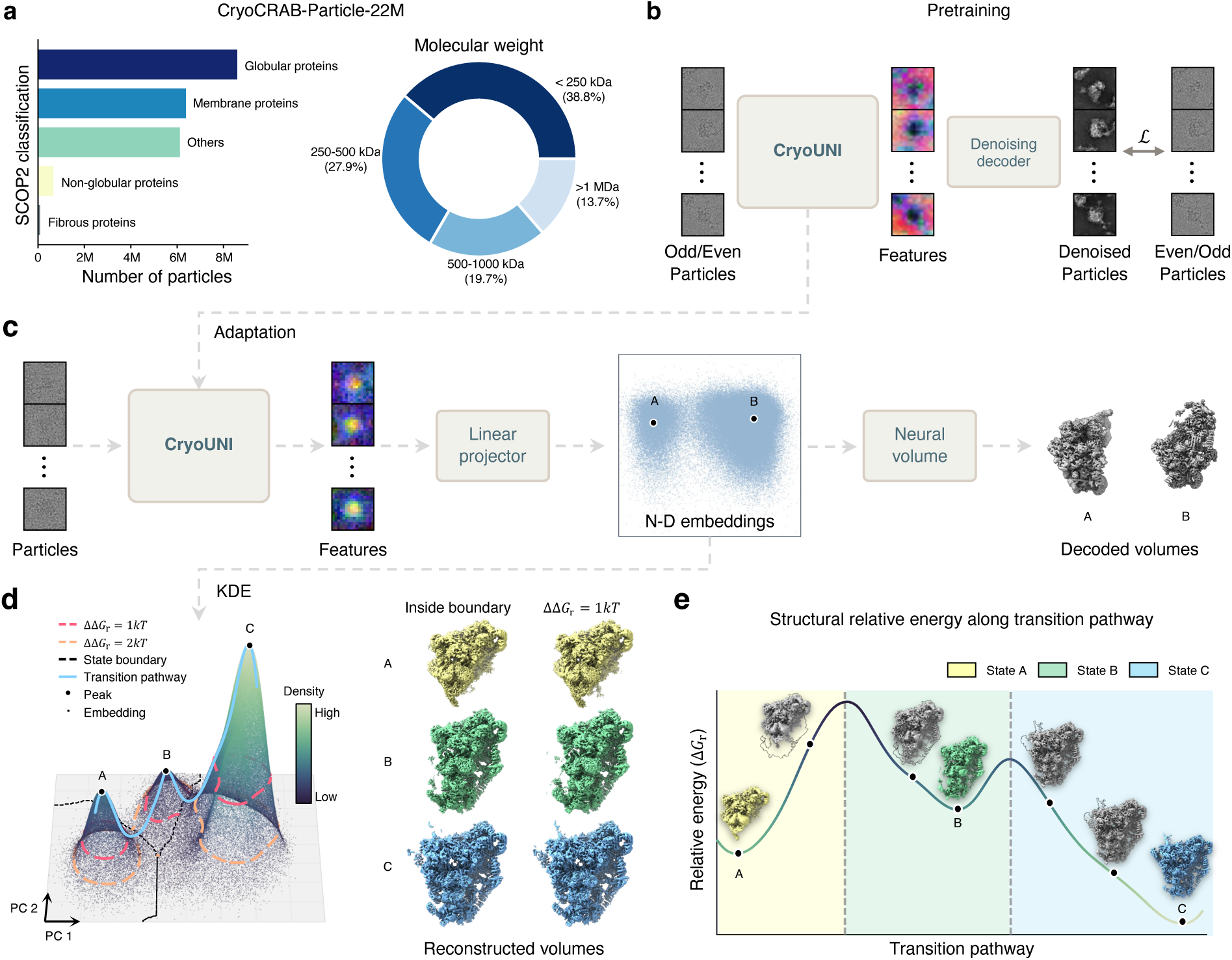
CryoUNI learns latent conformational landscapes from single-particle cryo-EM data. **a**, Composition of the CryoCRAB-Particle-22M pretraining dataset across SCOP2 structural classes and molecular weight ranges. **b**, Self-supervised pretraining of CryoUNI. Particles from one half-dataset are denoised using the complementary half as supervision, and we demonstrate particle images of 50S ribosomal intermediates [40]. **c**, Variational autoencoder architecture for downstream conformational analysis. A lightweight linear projector maps particle features into a low-dimensional embedding space, from which a neural volume decoder reconstructs the corresponding density volumes. **d**, WAVE (Watershed Analysis of Variational Embeddings) identifies discrete conformational states as local density maxima on the conformational landscape. Relative free energies are inferred from density ratios *via* Δ*G_r_* = *−k_B_T* ln(*ρ_A_/ρ_B_*). **e**, Continuous conformational trajectories traced between structural states, showing structural evolution and relative population of intermediates of yeast SSU processome assembly intermediates [41].

For downstream conformational analysis, the pretrained CryoUNI encoder is adapted to a target dataset *via* a variational autoencoder (VAE) architecture, mapping each particle image into a low-dimensional embedding space through a lightweight linear projector, from which a neural volume decodes the corresponding structural density (Fig. 1c). Notably, since the encoder is trained to suppress noise-driven variation, the density in the latent space is expected to reflect the underlying distribution of structural states, approximating a structural landscape that can be described by Boltzmann statistics.

To reveal a continuous structural landscape from such latent space, we first apply kernel density estimation (KDE) to the particle embeddings, followed by the watershed analysis of variational embeddings (WAVE) for automated state identification and transition pathway generation (Fig. 1d). Specifically, WAVE identifies local density maxima as discrete conformational states and draws their boundaries, revealing both dominant conformations and rare intermediates. The relative energy between states is then directly inferred from their density ratio *via* Boltzmann statistics, Δ*G_r_* = −*kT* ln(*ρ_A_/ρ_B_*), where *kT* denotes the thermal energy scale at the pre-freezing temperature. Continuous conformational pathways between states are traced along the landscape, revealing the structural transitions and relative population of intermediates (Fig. 1e). Together, CryoUNI reveals cryo-EM observations as a continuous probabilistic structural landscape, followed by WAVE analyses to reveal both discrete and continuous structural heterogeneity.

## 3 The latent space is physically grounded in the molecular conformational landscape

A central question for any learned conformational landscape is whether it reflects true physical reality. To test this, we compared the learned landscape with independent molecular dynamics (MD) simulations of integrin *αvβ*8 in extended-closed conformation [18]. This system exhibits continuous rotation of the flexible leg, which is not successfully resolved in the published density map. We reproduce such molecular motion through well-converged all-atom MD simulations comprising twenty independent 1-*µ*s trajectories (20 *µ*s aggregate; Fig. 2a). We then characterize it by mapping to a spherical coordinate system anchored at the head-leg hinge center of mass (COM), parameterizing leg orientation *via* polar (*θ*) and azimuthal (*ϕ*) angles (Fig. 2b). This coordinate system is applicable to both atomic models and reconstructed cryo-EM volumes, enabling direct comparison across representations. Projection of all sampled 160,000 MD snapshots onto this (*θ*, *ϕ*) space reveals that the leg motion is confined to a well-defined thermodynamic landscape (Fig. 2c and Supplementary Fig. C2a).

**Fig. 2.**
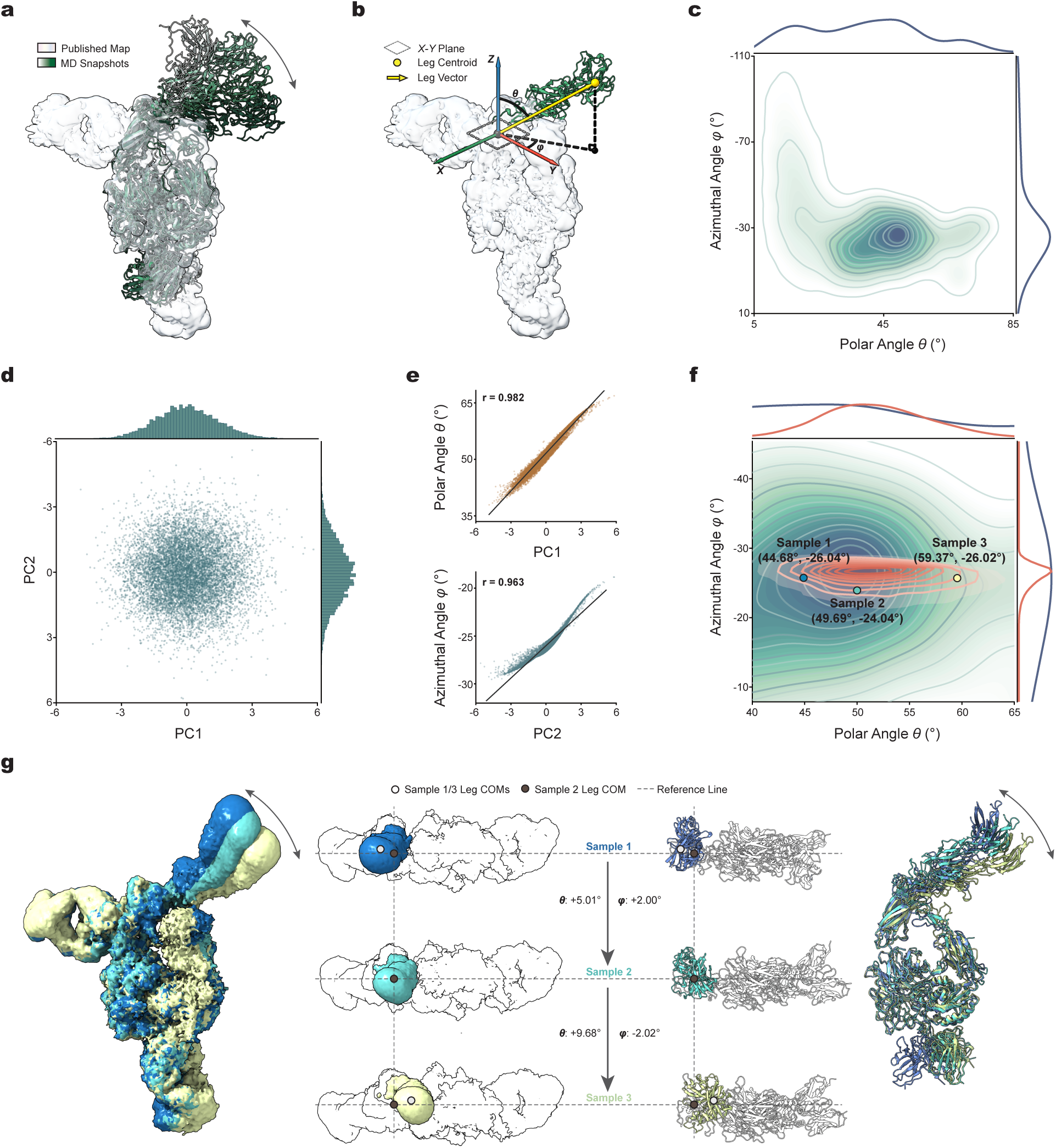
Physical grounding of CryoUNI latent conformational landscapes for integrin. *αvβ*8 **dynamics. a,** Conformational heterogeneity of the flexible leg of integrin *αvβ*8. Overlay of the published consensus cryo-EM map (EMPIAR-10345, grey surface) and representative MD snapshots (green cartoons). The distal leg density remains unresolved due to the continuous rotation captured in the MD ensemble. **b,** Unified spherical coordinate system for leg orientation. The origin is anchored at the head-leg hinge center of mass (COM); leg orientation is parameterized by polar angle *θ* and azimuthal angle *ϕ* (illustrated). A representative MD snapshot (green, non-leg regions omitted) and the cryo-EM map (grey surface) are shown. **c,** MD-derived thermodynamic conformational landscape. Two-dimensional probability density of (*θ, ϕ*) from 160,000 MD snapshots (twenty 1-*µ*s trajectories; 20 *µ*s aggregate). The leg motion is confined to a well-defined continuous region. **d,** Probabilistic latent space distribution of CryoUNI reconstructions. Principal component analysis of 10,000 decoded volumes reveals a continuous, unimodal distribution. **e,** Spontaneous disentanglement of physical coordinates. Pearson correlation between latent axes (PC1, PC2) and spherical coordinates (*θ, ϕ*). PC1 correlates with *θ* (*r* = 0.982); PC2 correlates with *ϕ* (*r* = 0.963). **f,** Projection of CryoUNI conformations onto the MD landscape. CryoUNI-derived conformations (orange contours) overlaid on the MD (*θ, ϕ*) probability density (green to navy). Three representative conformations are indicated. **g,** 3D structural validation of physical dynamics. CryoUNI density maps at three selected (*θ*, *ϕ*) coordinates (Sample 1-3: *θ* = 44.68°, 49.69°, 59.37°; *ϕ* = -26.04°, -24.04°, -26.02°), shown alongside their closest-matching MD atomic models. Leg orientations and local structural details are highly consistent across samples.

Applying CryoUNI to this dataset yields a latent space with a continuous, unimodal distribution (Fig. 2d). The learned latent axes align closely with the physical degrees of freedom captured by MD: PC1 correlates strongly with polar angle *θ* (*r* = 0.982), and PC2 with azimuthal angle *ϕ* (*r* = 0.963) (Fig. 2e). This level of correspondence, achieved without imposing biophysical priors, indicates that the latent representation captures the intrinsic conformational coordinates of the system and is therefore physically plausible.

We further evaluate this correspondence by projecting EM-reconstructed conformations into the MD-derived (*θ*, *ϕ*) landscape (Fig.2f). Coverage analysis based on highest-density regions (HDR) confirms that the MD conformational landscape encapsulates nearly all CryoUNI-generated samples (Supplementary Fig. C2b). As expected, cryo-EM captures the most populated structural states, while MD additionally further explores surrounding thermal fluctuations.

Finally, we selected three representatives (Sample 1–3) from the conformational landscape spanning the angular range of leg motion from the (*θ*, *ϕ*) distribution in Fig. 2f. For each sample point, we extract the corresponding CryoUNI-reconstructed density volume and the MD atomic snapshot with the closest (*θ*, *ϕ*) coordinates. The leg orientations observed in the CryoUNI density maps and the MD-derived structures are highly consistent (Fig.2g), as shown for three samples with polar angles *θ* of 44.68°, 49.69°, and 59.37°, and azimuthal angles *ϕ* of -26.04°, -24.04°, and -26.02°, respectively. As *θ* progressively increased, both CryoUNI and MD showed a gradual increase in leg swing amplitude. At each sample point, the rotation direction, amplitude, and local structural details all demonstrate high similarity between the CryoUNI reconstruction and the MD model. Moreover, the leg COM positions (labeled as Leg COMs in Fig.2g) aligned well with the corresponding (*θ*, *ϕ*) coordinates. CryoUNI not only recapitulates the global conformational landscape but also reconstructs density maps that faithfully mirror atomic-level dynamics (Supplementary Video X).

Together, these results provide evidence that the latent space of CryoUNI is not a mathematical artifact of dimensionality reduction but rather a deeply physically grounded conformational manifold. It decodes the intrinsic conformational degrees of freedom that govern the structural dynamics of integrin *αvβ*8.

## 4 Structural states and transitions emerge from the latent landscape

Having established the physical correspondence between the learned landscape and molecular conformational space, we next analyze the intrinsic structure of the latent landscape to resolve molecular heterogeneity (Fig. 1d,e). We introduce WAVE (Watershed Analysis of Variational Embeddings), which operates directly on the latent density to automatically identify conformational states as density peaks and reconstruct transitions by tracing high-density pathways, without imposing clustering assumptions or requiring a predefined number of states. This physical correspondence further defines a relative energy landscape described by Boltzmann statistics, in which high-density regions correspond to low-energy states.

We first evaluated WAVE on a simulated Ribosembly benchmark [19], which models the assembly of the 50S ribosomal subunit comprising 16 discrete compositional states. CryoUNI yields a well-organized latent landscape in which individual assemblies occupy compact, high-density regions (Extended Data Fig. 2a). WAVE automatically detects 16 regions as distinct density peaks and recovers all ground-truth states with high accuracy (99.4%). In contrast, other baselines achieve lower accuracy despite being provided with the correct number of clusters (Extended Data Fig. 2d), highlighting the importance of the quality of the latent representation itself. These results show that discrete compositional states emerge naturally as topological features of the latent landscape of CryoUNI. Beyond state identification, pairwise latent distances between density peaks strongly correlate with ground-truth structural similarity measured by normalized cross-correlation (Mantel test, *r* = −0.887, *p* = 0.001; Extended Data Fig. 2e). This further confirms that the latent landscape not only separates distinct states but also preserves their structural relationships, with neighboring peaks corresponding to structurally similar assemblies. The complete quantitative comparisons with additional baselines are provided in Extended Data Table 1.

To assess scalability to larger compositional mixtures, we evaluated the Tomotwin-100 dataset, which contains 100 distinct structural classes spanning a broad range of molecular weights. CryoUNI produces a latent landscape in which WAVE automatically detects all 100 density peaks (Extended Data Fig. 2f). Quantitatively, CryoUNI achieves near-perfect classification accuracy regardless of the clustering algorithm used (K-means: 99.98%, GMM: 99.97%, WAVE: 99.96%), substantially outperforming all other methods (Extended Data Fig. 2g; Extended Data Table 1). In contrast, applying WAVE to latent spaces learned by other methods leads to substantially degraded performance (Extended Data Table 1), suggesting that their latent density topology does not faithfully reflect the underlying compositional structure.

Finally, we evaluated a dataset dominated by continuous conformational variability (IgG-1D), where discrete state classification is no longer applicable. The CryoUNI latent landscape captures the underlying conformational continuum as a continuous manifold, with WAVE tracing a closed trajectory that recovers the full conformational cycle (Extended Data Fig. 3a,b). To quantitatively assess reconstruction quality, we measured per-particle AUC-FSC, which evaluates how faithfully each particle’s structure is recovered. CryoUNI achieves the highest mean AUC-FSC (0.368) with the smallest variance across all methods, significantly outperforming existing approaches (Extended Data Fig. 3c).

Together, these results demonstrate that structural states and transitions emerge directly from the density and topology of the CryoUNI latent landscape, providing a unified representation across discrete compositional mixtures and continuous conformational dynamics.

## 5 The latent landscape reveals previously unresolved intermediate states

We next ask whether the learned landscape can reveal previously unresolved states in experimental data. We analyzed a cryo-EM dataset of LIS1-mediated dynein remodeling under ATP conditions [10], which captures the LIS1-dynein activation process. In the initial latent landscape, particles organize into well-separated high-density basins corresponding to the three major structural classes along the activation pathway: open-bent (State A), open-straight (State B), and motor-bound (State C) (Fig. 3a). The relative energy profile along the WAVE-traced transition pathway reveals these states as distinct energy minima separated by barriers, outlining the global organization of the activation pathway.

**Fig. 3.**
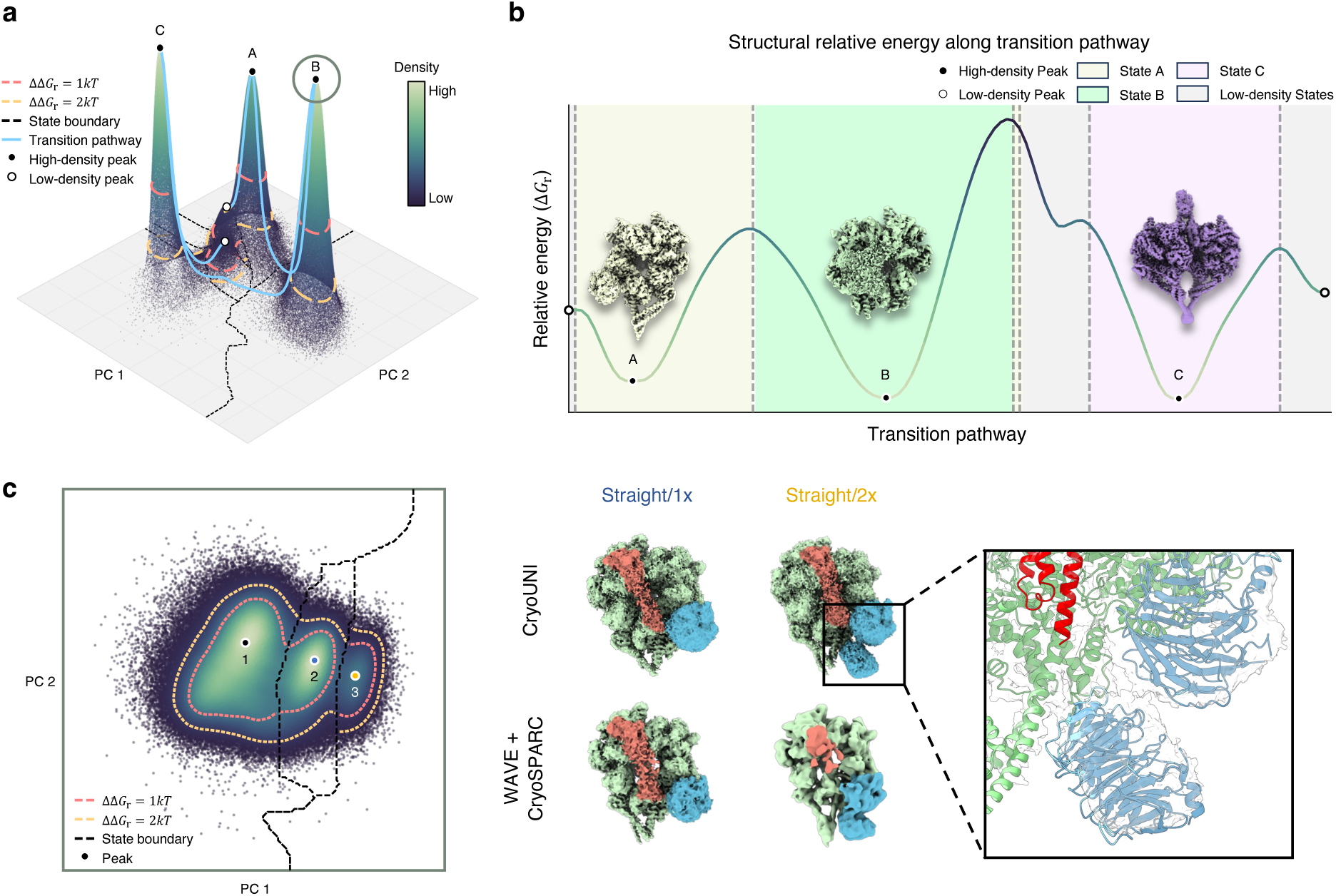
Hierarchical CryoUNI analysis resolves the dynein activation pathway and novel binding mode from EMPIAR-12715. a, Global latent space representation of the EMPIAR-12715 dataset projected onto the first two principal components (PC 1 and PC 2). **b,** The structural relative energy along the transition pathway in **a**. The color gradient indicates particle density, revealing well-separated regions corresponding to major structural classes such as open-bent (State A), open-straight (State B), and motor-bound (State C) configurations. **c,** Hierarchical analysis of local density structures within open-straight region. Comparisons between the CryoUNI and WAVE + CryoSPARC frameworks are shown. Relative energy boundaries (ΔΔ*Gr* at 1 *kT* and 2 *kT*) delineate structurally distinct basins. This progressive refinement resolves finer variability, including distinct LIS1 stoichiometries (1*×*LIS1 and 2*×*LIS1) and a newly found straight sub-state featuring a second LIS1-associated density. Structures are colored by subunit: Dynein linker (Warm Terracotta), Dynein motor (Pale Celadon), and LIS1 (Vibrant Ocean).

Within each major structural class, the landscape further reveals hierarchical organization, with multiple density peaks indicating the presence of sub-states (Extended Data Fig. 4a). We focus here on the open-straight region, where this hierarchical analysis uncovers previously unresolved heterogeneity. The local landscape reveals multiple density peaks with distinct occupancies (Fig. 3b). Particle subsets associated with these peaks yield structurally distinct reconstructions that differ in their LIS1 binding mode. One population (Straight/1x) contains a single LIS1 copy. Notably, another population (Straight/2x)—previously unidentified—shows an additional density feature suggesting a second LIS1 molecule bound at the dynein stalk-buttress interface (Fig. 3c). This low-population state, characterized by a second LIS1 binding event, provides a new structural basis for the stepwise activation mechanism of dynein [11]. These results demonstrate that the latent landscape can resolve low-population intermediates defined by distinct binding modes, which conventional methods merge or overlook.

Extending this analysis across all three structural classes reveals a total of 12 sub-states from the latent landscape (Extended Data Fig. 4a,b). Within the motor-bound class, the landscape recovers the autoinhibited *ϕ* state and early intermediate conformations consistent with previously reported structures [10]. The open-bent class resolves multiple sub-states with distinct LIS1 binding modes, including variations in copy number and binding site that are not successfully separated using conventional approaches. All identified sub-states yield structurally consistent reconstructions at resolutions ranging from 2.75 to 6.44 Å (Extended Data Fig. 4b), confirming that density peaks correspond to genuine conformational populations. Several states achieve improved resolution compared to the original analysis, indicating that landscape-guided particle selection produces more homogeneous subsets. Every previously reported conformation is recovered, together with a new Straight/2x state that has been resolved.

Together, these results demonstrate that the latent landscape unlocks the capability for hierarchical analysis of molecular heterogeneity, from the coarse separation of dominant structural classes to the fine-grained resolution of low-population intermediates defined by distinct binding modes.

## 6 The latent landscape enables energy-guided reconstruction and pathway generation

Finally, we ask whether the conformational landscape can improve the reconstruction quality. In this conformational landscape described by Boltzmann statistics, density ratios directly infer yield relative energies, making high-density regions a principled basis for energy-guided particle selection. We apply this principle to a cryo-EM dataset of the KCTD5/CUL3^NTD^/G*βγ* complex [12], in which four discrete conformations have been previously resolved, but continuous dynamics were previously accessible only through morphing interpolations. In the latent landscape, particles organize into four well-separated high-density basins corresponding to the reported structural states (Fig. 4a). WAVE automatically identifies these four states with high agreement to the original classification (Fig. 4c), with over 91% of particles in each class correctly assigned.

**Fig. 4.**
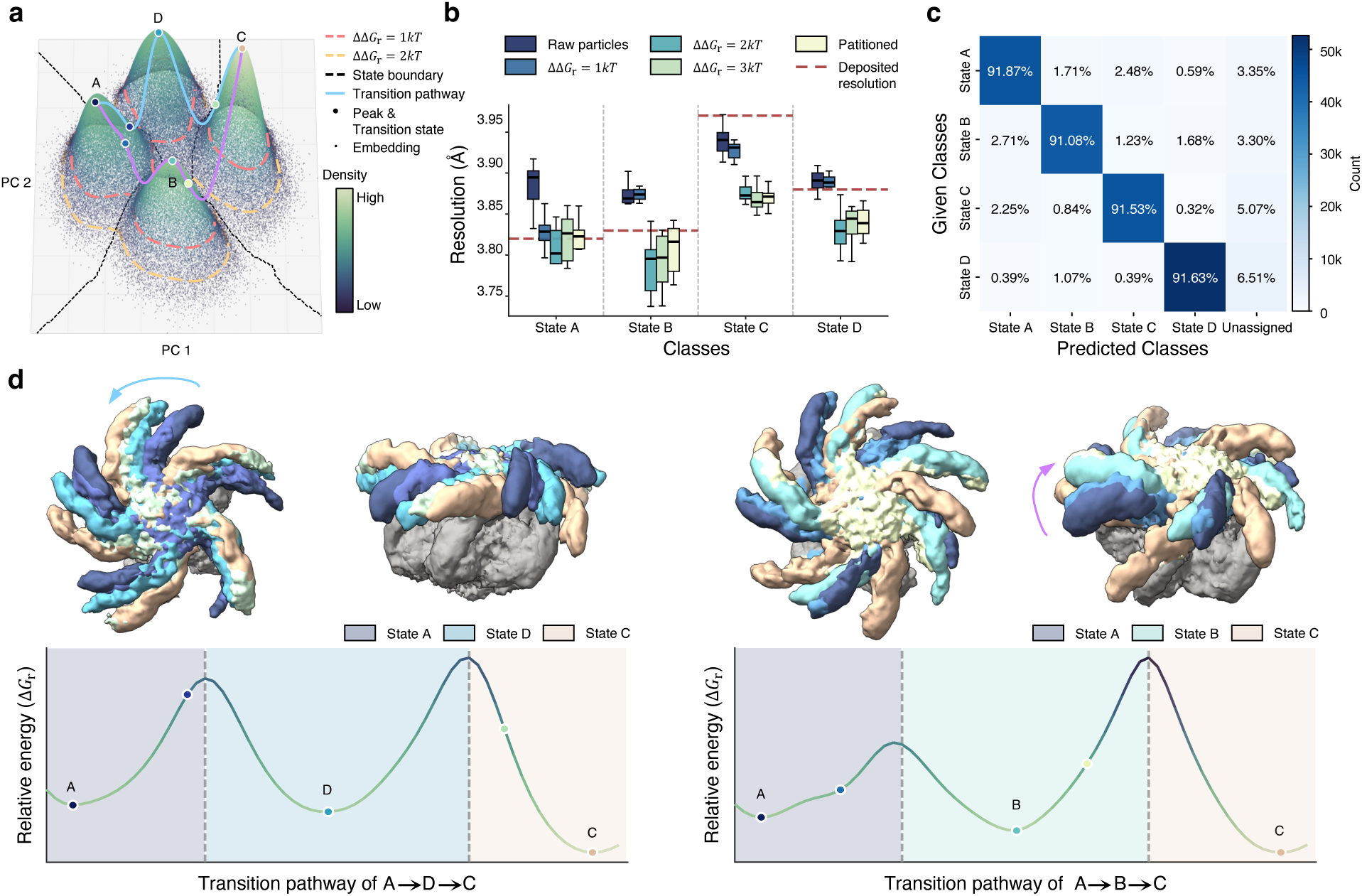
Energy-guided reconstruction and conformational trajectory analysis of the KCTD5/CUL3^NTD^/G*βγ* complex. **a**, CryoUNI latent landscape colored by density, with four detected peaks (A–D) corresponding to the major conformational states. Dashed lines indicate watershed boundaries (black), and relative energy contours at ΔΔ*G*r = 1 *kT* (red) and 2 *kT* (orange). Trajectories traced by WAVE are shown in purple. **b**, Reconstruction resolution as a function of energy-guided particle selection. For each state, particle subsets are selected at increasing ΔΔ*Gr* thresholds (1*kT*, 2*kT*, 3*kT*) and at the watershed boundary (*n*_State A_ = 48,977; *n*_State B_ = 48,890; *n*_State C_ = 57,006; *n*_State D_ = 47,975 total particles before threshold filtering). Box plots show the median (center line), interquartile range (IQR, box edges, 25th–75th percentiles), and 1.5*×* IQR (whiskers); outliers are plotted individually. Red dashed lines indicate the deposited resolution from the original analysis. Resolution was estimated from *n* = 10 independent half-map splits per condition. Intermediate thresholds yield improved resolution across all states. **c**, Confusion matrix comparing WAVE-identified states with the original classification. Diagonal entries show agreement exceeding 91% for all four states without prior specification of state number. **d**, Continuous conformational pathways connecting the four states. Left: trajectory A*→*D*→*C with representative volumes colored by state identity (State A, State D, State C) and the corresponding relative energy profile (Δ*Gr*) along the pathway. Right: trajectory A*→*B*→*C with representative volumes and energy profile. Energy barriers between states are directly resolved from the latent density.

With the four states resolved, we next leverage the energy structure of the landscape for particle selection. For each state, we select particle subsets at increasing relative energy thresholds (ΔΔ*G*_r_) derived from latent density (Fig. 4a). Across all states, energy-guided selection improves both reconstruction consistency and resolution compared to using the full particle set (Fig. 4b). Subsets at intermediate ΔΔ*G*_r_ thresholds achieve the best performance, balancing the removal of structurally heterogeneous particles with retention of sufficient signal. These results demonstrate that latent density provides a principled criterion for particle selection without heuristic filtering.

The landscape also captures conformational transitions between the resolved states. The four density basins are connected by continuous pathways, and tracing these pathways identifies two dominant conformational trajectories: A→D→C and A→B→C (Fig. 4d). This experimental resolution of the trajectory deepens our understanding of its substrate recognition and ubiquitination at the specific lysine site (K23) on the G*βγ* substrate across various reactive states [20]. The relative energy profiles along each trajectory reveal the energy barriers separating intermediate states. These trajectories recover the principal motions previously inferred from morphing interpolation between known states [12], providing direct experimental resolution of continuous conformational dynamics for this complex from cryo-EM data alone.

Together, these results demonstrate that a single physically grounded landscape enables both high-resolution structure determination through energy-guided particle selection and direct recovery of transition pathways from cryo-EM data.

## 7 Discussion

We have realized cryo-EM as structural landscape microscopy, in which latent density reflects the probabilistic distribution of conformational states and defines a relative energy landscape described by Boltzmann statistics. This representation unifies the analysis of discrete compositional states, continuous conformational dynamics, and energy-guided particle selection within a single physically grounded framework.

The key conceptual shift is from treating the latent space as a computational intermediate for volume generation to interpreting it as a conformational landscape grounded in physical terms. That said, previous methods have made substantial progress in modeling structural heterogeneity from cryo-EM data. Discrete classification approaches [1, 2] resolve major conformational states but require specifying the number of classes and cannot capture continuous dynamics. Continuous methods such as cryoDRGN [3], 3DVA [4], 3D Flex [5], and RECOVAR [7] represent variability in learned latent spaces, enabling volume generation along continuous pathways. More recently, foundation-model-based approaches [8, 21] leverage learned priors to improve reconstruction quality. While these methods have advanced heterogeneous reconstruction, the physical interpretability of the latent space itself has received less attention. Our work builds on these advances by showing that, with a latent structural landscape, density and distance directly encode state occupancy, structural relationships, and transition pathways. This establishes a physically grounded basis for downstream analysis.

The interpretation of latent density as a relative energy connects cryo-EM analysis to statistical mechanics, but several limitations remain. First, the observed state distribution may deviate from true equilibrium occupancy due to preferred orientations, uneven ice thickness, and the particle curation procedure. Second, the latent representation is learned from noisy 2D projection images, in which undesired features such as membrane and detergent densities may dominate over internal conformational changes. For such systems, incorporating structural priors could help disentangle biologically meaningful variation from environmental contributions. Last, the latent density has not been directly linked to a physical energy function, which is why we report relative rather than absolute energy; incorporating physics-informed constraints could help establish this connection.

More broadly, this work suggests a broader shift in how cryo-EM data are interpreted: from determining individual structures to characterizing conformational landscapes. Several directions follow naturally. Combining learned conformational landscapes with physics-based simulations may enable quantitative free energy estimation directly anchored in experimental observation, building on the connection between cryo-EM and molecular dynamics established here. Extending the approach to in situ cryo-electron tomography [22] data could enable characterization of conformational landscapes within their native cellular context. Finally, integrating learned conformational landscapes with complementary biophysical and computational approaches may ultimately connect structural descriptions with the kinetic and thermodynamic principles that govern molecular function.

## Methods

### Large-scale pretraining

#### Large-scale particle dataset

We constructed a large-scale particle image dataset, CryoCRAB-Particle-22M, to train and evaluate CryoUNI. The dataset comprises 22,631,283 particle images extracted from 173,997 micrographs spanning 746 distinct protein species, built upon the CryoCRAB dataset [23]. Particles were extracted using an iteratively optimized scheme: initial picking by particle picking adapted cryoUNI (Extended Data Fig. 1c) was followed by two rounds of 2D classification (200 and 50 classes, respectively), and this picking-curating-retraining cycle was repeated for six iterations to yield a high-purity particle collection. All particles are stored in HDF5 format with float16 precision at their native resolution. To support self-supervised pretraining, each particle is stored as a full-difference pair: the full image is generated by averaging all raw movie frames, while odd and even frame averages provide independent half-images whose subtraction yields a pure noise estimate. During training, geometric augmentation is applied, including random horizontal flipping and random rotation over {0^◦^, 90^◦^, 180^◦^, 270^◦^}. All images are preprocessed with CTF phase-flipping, contrast normalization, and Z-score standardization following the CryoCRAB protocol [23].

#### Architecture of CryoUNI

Our model builds upon the Vision Transformer (ViT) architecture, including modern design choices from recent Transformer models [24] (Extended Data Fig. 1a). Given an input image *x* ∈ R*^H^*^×*W*^, we first divide it into *H/P* × *W/P* non-overlapping patches with patch size *P* = 16. All patches are linearly projected into *D*-dimensional embeddings and concatenated with one class token and four register tokens [25] to form the input sequence. This sequence is fed into a stack of *L* Transformer blocks. Each block consists of a multihead self-attention mechanism with 2D Rotary Position Embedding (RoPE) [26], followed by a SwiGLU feed-forward network. Residual connections and RMSNorm are applied around each sub-layer for training stability. We use two configurations: small (*D* = 384, *L* = 12) and base (*D* = 768, *L* = 12).

#### Denoising-reconstruction pretraining

In the first pretraining stage, we adopt the denoising-reconstruction pretraining strategy from DRACO [17], using odd-even-full triplets generated from the CryoCRAB full-diff pairs as training samples. Take the encoded feature as input, a transformer-based decoder is employed to reconstruct the original image; it projects the *D*-dimensional encoder features to 576 dimensions and refines them through 8 Transformer blocks. Each patch token is then decoded into a *P* × *P* pixel patch. To accommodate the variable resolution of particle images without information loss from rescaling, we follow the approach of Dehghani et al. [27], dynamically converting images into token sequences at their native resolution. This decoder is discarded after pretraining. The pretraining combines masked image modeling loss on masked regions [28] (against the high-SNR full image) with Noise-to-Noise loss on visible regions (against the paired half-image), as described in [17]. To align with downstream tasks, input particle images are CTF-corrected, while the corresponding CTF modulates the decoder output before computing the loss against the original non-CTF-corrected reference images. The combined loss function is formally defined as:

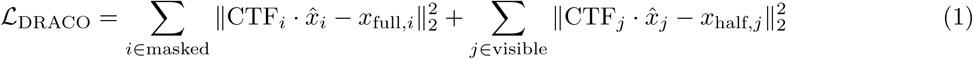

where *x*^ represents the decoder output. We use the AdamW optimizer with a base learning rate of 1.5×10^−4^, weight decay of 0.05, and base EMA rate of 0.995. The image mask ratio is set to 0.75. The model is trained for 800 epochs with a batch size of 4096, including a learning rate warmup for the first 40 epochs.

#### Denoising pretraining

In the second pretraining stage, we further enhance the model’s denoising capability to capture fine-grained structural details. We leverage odd-even image pairs based on the Noise-to-Noise principle [16] and integrate CryoUNI with the Dense Prediction Transformer (DPT) [29] framework to construct a multi-scale denoising network. Feature maps are extracted from CryoUNI at every Transformer block and transformed into a multi-scale feature pyramid, which is progressively upsampled and fused through cascaded interpolation and residual convolution. To preserve frequency-domain integrity, a two-layer convolutional network preprocesses the input image and adds it to the final fused features before producing the denoised output through another two-layer convolutional network (Extended Data Fig. 1d). The Noise-to-Noise objective is optimized using the expected loss between the predicted output from one half-image and its paired half-image:

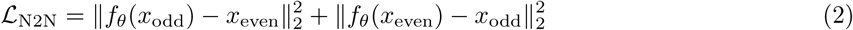

where *f_θ_*denotes the denoising network. To minimize this loss, we employ the AdamW optimizer with a base learning rate of 1.0e-4, a weight decay of 0.05, and a base exponential moving average (EMA) rate of 0.99. The batch size is 2048, and the model is trained for a total of 50 epochs, with a learning rate warmup applied for the first 5 epochs.

### Heterogeneous reconstruction

#### Model architecture

To resolve distinct conformational states of proteins from noisy particle images, we developed a heterogeneity reconstruction framework based on a Variational Autoencoder (VAE). This framework adopts the fundamental architecture of cryoDRGN [3] with optimizations in both the encoder and decoder components (Extened Data Fig. 1d). On the encoder side, we integrated a pretrained CryoUNI for end-to-end fine-tuning; it extracts class and patch tokens from images and employs a linear neck to aggregate local features, which are subsequently mapped to a latent space to generate embeddings **z** representing conformational heterogeneity. For the decoder, we implemented a volume decoder based on implicit neural representations. Taking 3D coordinates and embeddings **z** as inputs, the decoder performs high-frequency encoding of coordinates via random Fourier features with 384 default Fourier embedding channels. The intensities in the Hartley domain are predicted using a 3-layer SwiGLU network with 384 embedding channels and residual connection and LayerNorm.

#### Adaptation

The training of the heterogeneous reconstruction model was optimized using the AdamW optimizer. We set the base learning rate to 1 × 10^−4^, which was scheduled following a cosine decay strategy. Training was performed on a single NVIDIA A40 GPU with a total batch size of 32 and a default training epochs of 50. To balance reconstruction fidelity with latent space regularization, the VAE objective function was weighted:

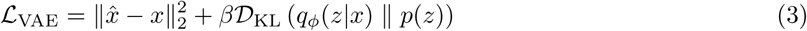

where L_VAE_ measures the data likelihood (reconstruction error in the Hartley domain), D_KL_ is the Kullback-Leibler divergence between the approximate posterior *q_ϕ_*(*z*|*x*) and the prior *p*(*z*), and the coefficient *β* is specifically set to 0.125.

### Watershed analysis of variational embeddings (WAVE)

#### Density estimation and peak detection

To characterize the conformational landscape, we project CryoUNI features into a *D*-dimensional latent space (*D* ∈ {2, 3, 4}) via principal component analysis (PCA), yielding a set of latent coordinates {**z***_i_*}*^N^*.

Similar to RECOVAR [7], a continuous probability density field is then estimated using kernel density estimation (KDE):

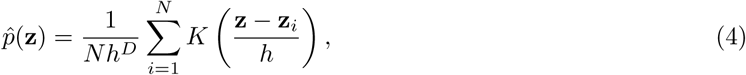

where *K*(·) is an isotropic Gaussian kernel and the bandwidth *h* is determined by Scott’s rule. The density field is evaluated on a uniform grid with resolution 256^2^ (*D* = 2), 128^3^ (*D* = 3), or 64^4^ (*D* = 4). Density peaks are identified as local density maxima using a multi-dimensional maximum filter with a 4*^D^*neighborhood. To suppress noise-induced spurious peaks, only maxima with density at least 5% of the global maximum are retained.

#### Transition pathway analysis

To trace continuous transition pathways between structural states, we adopt a coarse-to-fine strategy. First, a Traveling Salesman Problem (TSP) solver determines the global ordering of density peaks by minimizing the total path length in latent space, ensuring that the traversal follows physically plausible transitions through high-density regions. Local pathways between adjacent peaks are then refined by solving the Eikonal equation:

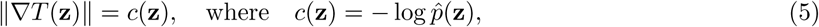

where *T* (**z**) represents the cumulative transition cost. The cost function *c*(**z**) is the negative log-density, consistent with the potential energy under Boltzmann statistics, guiding pathways through the highest-density regions of the landscape. We solve this equation on the previously defined grid using the Fast Marching Method (FMM) to extract the most probable transition pathway between states.

#### Partitioning and particle selection

To define the boundaries of structural states, we apply a marker-controlled watershed algorithm using the detected density peaks as initial markers. This partitions the latent space into non-overlapping domains, and each particle is assigned to a domain based on its latent coordinate **z***_k_*. To enhance conformational purity within each domain, we apply an energy-based filtering criterion. For the *i*-th state, only particles satisfying the following condition are retained:

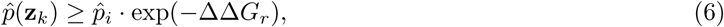

where *p̂_i_* is the peak density of the *i*-th conformational center and ΔΔ*G_r_* is the relative energy threshold. This selects particles within a defined energy range of each density peak.

### Metrics

#### Resolution estimation via Fourier shell correlation (FSC)

The resolution of cryo-EM reconstructions was estimated using the Fourier Shell Correlation (FSC) criterion [30]. The FSC measures the normalized cross-correlation between two independently reconstructed half-maps as a function of spatial frequency. For a given spatial frequency shell *k*, the FSC is defined as:

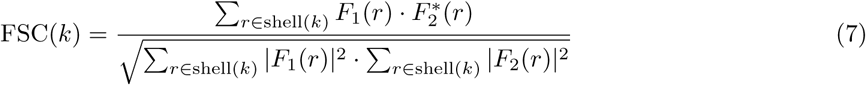

where *F*_1_(*r*) and *F*_2_(*r*) are the complex structure factors of the two half-maps at spatial frequency vector *r*, and *F*_2_^∗^(*r*) denotes the complex conjugate of *F*_2_(*r*). The overall resolution was determined at the gold-standard FSC = 0.143 threshold [31].

To provide a more comprehensive measure of global structural fidelity, we additionally computed the Area Under the Curve of the FSC (AUC-FSC) proposed by Jeon et al. [19]:

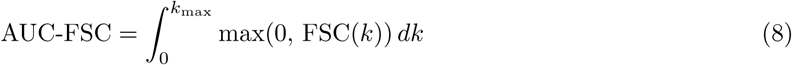

where *k*_max_ is the Nyquist frequency. By integrating the correlation over all spatial frequency shells, this metric captures the cumulative signal quality across all resolution bands, rather than relying on a single-threshold estimate. To account for the stochastic effects of random particle half-set splitting, we performed multiple independent reconstruction trials for each state and report the mean and variance of the resulting resolution estimates.

#### Normalized cross-correlation (NCC) and Mantel test

To assess whether the geometric organization of the CryoUNI latent landscape faithfully preserves the physical relationships between conformational states, we evaluated the correlation between pairwise latent distances and 3D structural similarities. The structural similarity between ground-truth volumes was quantified using the Normalized Cross-Correlation (NCC) score:

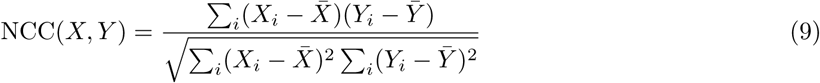

where *X_i_* and *Y_i_* represent the voxel intensities of two aligned 3D volumes, and *X̄* and *Ȳ* are their respective mean voxel intensities. The pairwise distance between states in the latent representation was computed as the Euclidean distance between their corresponding detected density peaks:

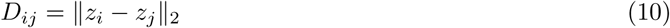

The correspondence between the latent topology and true structural variance was evaluated using the Mantel test [32], which assesses the correlation between the pairwise latent distance matrix and the NCC similarity matrix while accounting for the non-independence of distance matrix elements. The test yields the Pearson correlation coefficient *r* and a permutation-based *p*-value (*P*_Mantel_).

#### Clustering accuracy

To quantitatively evaluate the ability of the learned latent representations to disentangle distinct conformational states, we measured the clustering accuracy by comparing predicted cluster labels against ground-truth class labels. The clustering accuracy is defined as the maximum proportion of correctly assigned particles over all possible label permutations between the predicted labels *ŷ_i_* and the ground-truth labels *y_i_*:

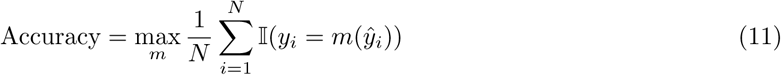

where *N* is the total number of particles, I(·) is the indicator function, and *m* is a permutation mapping that assigns each predicted cluster to a ground-truth class. The optimal mapping *m* was solved using the Hungarian algorithm.

#### Adjusted Rand Index (ARI) and Adjusted Mutual Information (AMI)

To evaluate clustering quality on the compositional heterogeneity benchmarks (Ribosembly and Tomotwin-100), we additionally report the Adjusted Rand Index (ARI) and Adjusted Mutual Information (AMI). Both metrics compare predicted cluster assignments against ground-truth labels and are corrected for chance, yielding a value of 0 for random labelling and 1 for perfect agreement. ARI measures the fraction of particle pairs that are consistently co-clustered or separated, adjusted by the expected value under a random permutation model [33]. AMI quantifies the mutual information between the two partitions, normalized by the entropy of the marginals and adjusted for chance agreement [34].

### Statistics and reproducibility

All statistical tests reported in this study are two-sided. No statistical method was used to predetermine sample sizes; particle counts are determined by the experimental datasets deposited in EMPIAR and the simulated benchmarks from CryoBench. No randomization or blinding was applicable to this computational study. No data were excluded from analyses except as described in the Data section, where 2D classification was applied to remove non-particle images (ice contamination, false positives) following standard cryo-EM data processing procedures.

The Mantel test [32] was used to evaluate the correspondence between pairwise latent distances and ground-truth structural similarities (NCC) on the Ribosembly dataset (*n* = 16 states, 120 unique pairwise comparisons; Extended Data Fig. 2e). Statistical significance was assessed using 999 random permutations of distance matrix rows and columns (Pearson *r* = −0.887, *p <* 0.001).

Pearson correlation coefficients were computed between CryoUNI latent principal components and the MD-derived spherical coordinates (*θ* and *ϕ*) for the integrin *αvβ*8 dataset (*n* = 10,000 decoded volumes; PC1 vs. *θ*: *r* = 0.982; PC2 vs. *ϕ*: *r* = 0.963; Fig. 2e).

For the IgG-1D continuous heterogeneity benchmark, per-conformation AUC-FSC was evaluated by stratified sampling: 10 particles were randomly drawn from each of the 100 conformational classes, yielding *n* = 1,000 reconstructed volumes per method. The AUC-FSC was computed for each volume against its corresponding ground-truth density, and results are reported as mean ± s.d. (Extended Data Fig. 3c). CryoUNI achieved the highest mean AUC-FSC (0.368) with the smallest variance. Pairwise comparisons between CryoUNI and each baseline method were performed using the two-sided Mann–Whitney *U* test on the 1,000 AUC-FSC values (CryoUNI vs. CryoDRGN: *U* = 998,910, *P <* 10^−300^; CryoUNI vs. 3DVA: *U* = 999,000, *P <* 10^−300^; CryoUNI vs. RECOVAR: *U* = 999,003, *P <* 10^−300^; CryoUNI vs. OPUS-DSD: *U* = 999,592, *P <* 10^−300^; CryoUNI vs. CryoDECO: *U* = 999,000, *P <* 10^−300^; CryoUNI vs. 3D Flex: *U* = 937,913, *P* ≈ 4.4 × 10^−252^). The near-maximal *U* statistics (theoretical maximum *U*_max_ = 10^6^ for *n*_1_ = *n*_2_ = 1,000) indicate that CryoUNI’s AUC-FSC values exceeded those of every baseline in 93.8–99.96% of all pairwise comparisons, reflecting near-complete distributional separation between CryoUNI and each competitor.

Reconstruction resolution was determined using the gold-standard FSC = 0.143 criterion [31] computed from independently refined half-maps.

In Supplementary Fig. C2a, error bars represent mean ± s.d. of the overlap between sub-sampled and full MD landscapes, computed from 100 random subsets of *k* trajectories drawn from the 20 independent 1-*µ*s MD simulations.

For the KCTD5/CUL3^NTD^/G*βγ* analysis (Fig. 4c), WAVE classification agreement with the original published states was quantified as the percentage of particles correctly assigned per class (*>*91% for all four states; *n* = 202,484 total particles).

Molecular dynamics simulations comprised 20 independent 1-*µ*s trajectories initiated from the same starting model with different random initial velocities (20 *µ*s aggregate). Only the 200–1,000 ns portion of each trajectory was used for quantitative analysis, yielding 160,000 snapshots at 0.1 ns intervals.

### Data

#### Integrin αvβ8 complex

We analyzed the EMPIAR-10345 dataset [18], which captures the integrin *αvβ*8 bound to latent transforming growth factor-*β* (L-TGF-*β*) and inhibitory antibody fragments. From 1,577 micrographs (after excluding 67 low-quality images), we extracted 559,086 particles using the CryoUNI particle picker (confidence threshold 0.0001) and downsampled to a box size of 128 pixels (3.152 Å/pix). After 2D classification in cryoSPARC, 110,148 clean particles were retained. We performed iterative refinement through three rounds of CryoUNI heterogeneous reconstruction (*z* = 8) with 10-component GMM clustering, progressively removing poorly resolved particles and resolving two compositional states (Fab-bound and unbound). After excluding Fab-lacking particles, a compositionally homogeneous stack of 57,546 Fab-bound particles was re-extracted at the unbinned pixel size (1.345 Å/pix, box size 300) and subjected to final non-uniform refinement (3.97 Å). A third round of heterogeneous reconstruction (*z* = 8) was performed on this stack for the final conformational landscape analysis.

#### KCTD5/CUL3^NTD^/Gβγ complex

We analyzed the EMPIAR-11734 dataset [12] to resolve the conformational landscape of the KCTD5/CUL3^NTD^/G*βγ* complex. We directly imported the curated particle stacks from the original study, yielding 202,484 particles (box size 324, pixel size 1.03 Å/pix) previously assigned to four states: State A (48,977), State B (48,890), State C (57,006), and State D (47,975). CryoUNI heterogeneous reconstruction was trained for 50 epochs with an 8-dimensional latent space (*z* = 8). WAVE partitioned the latent landscape into four conformational basins consistent with the original classification. To resolve continuous transitions between states, we performed PCA on the latent embeddings and extracted transitional volumes along WAVE-traced pathways through the principal component subspace.

#### LIS1-mediated dynein activation

We analyzed the EMPIAR-12715 dataset [10] to resolve the conformational landscape of LIS1-mediated dynein remodeling. From 12,753 usable micrographs, the CryoUNI particle picker (confidence threshold 0.0001) identified 4,943,683 particles. After two rounds of 2D classification in cryoSPARC, 965,465 particles were retained. Ab initio reconstruction (6 classes) followed by heterogeneous refinement separated the dataset into three major structural classes: open-bent (230,620 particles), open-straight (346,948 particles), and motor-bound (143,010 particles). Particles were re-extracted at 0.935 Å/pix (box size 352) and refined to 2.64, 3.03, and 2.71 Å, respectively.

To construct the global conformational landscape, poses from the three classes were aligned using cryoSPARC Align 3D, and the combined particle set was used to train a global CryoUNI reconstruction model for 50 epochs. Using WAVE, the global latent landscape recovered the three major classes with refined particle distributions: open-bent (165,014 particles), open-straight (346,896 particles), and motor-bound (208,668 particles). To resolve sub-states within each class, the three subsets were processed independently with CryoUNI for 50 epochs, generating local conformational landscapes. Latent dimensions were optimized per class (*z* = 8 for motor-bound, *z* = 4 for open-bent and open-straight). WAVE partitioned each local landscape into sub-states, and the corresponding particle subsets were re-imported into cryoSPARC for final high-resolution reconstruction.

#### CryoBench datasets

We evaluate CryoUNI on three simulated datasets from the CryoBench benchmark [19], each targeting a distinct form of structural heterogeneity. **Ribosembly** comprises 335,240 particle images (128×128, 6 Å/pix, SNR 0.01) modeling the stepwise assembly of the 50S ribosomal subunit across 16 discrete compositional states, providing a benchmark for state identification and latent space organization. **Tomotwin-100** contains 100,000 particle images (128×128, 9 Å/pix, SNR 0.01) spanning 100 distinct structural classes with a broad range of molecular weights, testing scalability to high-complexity compositional mixtures. **IgG-1D** contains 100,000 particle images (128×128, 6 Å/pix, SNR 0.01) simulating continuous conformational variability of an IgG antibody along a single degree of freedom, assessing reconstruction fidelity across the conformational landscape.

For Ribosembly, we train CryoUNI for 25 epochs with latent dimension *D* = 4. For Tomotwin-100, we train CryoUNI for 100 epochs with latent dimension *D* = 8. For IgG-1D, we train CryoUNI for 20 epochs with latent dimension *D* = 8. All models use a batch size of 32 and the AdamW optimizer with a learning rate of 1 × 10^−4^ and KL divergence weight of 0.0125. All experiments were conducted on a single NVIDIA A40 48G GPU. We compare against cryoDRGN [3], 3DVA [4], RECOVAR [7], OPUS-DSD [6], CryoDECO [8], and 3D Flex [5] as baselines. Detailed configurations for each baseline are provided in the Baselines section below.

### Baselines

All methods were run using their official implementations with default hyperparameters unless otherwise noted. Latent embeddings from each method were used for downstream clustering (K-means, GMM) and WAVE analysis.

#### CryoDRGN

CryoDRGN [3] models continuous structural heterogeneity using an amortized variational autoencoder. Each particle image is encoded into a low-dimensional latent variable, from which a coordinate-based neural network decodes 3D density volumes in Fourier space. We used the official CryoDRGN implementation (version 3.4.4) with default network architecture. The model was trained for 25 epochs with a latent dimension of 8 on each dataset. Image poses were fixed to the consensus values estimated by cryoSPARC.

#### 3DVA

3D Variability Analysis (3DVA) [4] fits a linear subspace model to cryo-EM data, representing conformational variability as a set of orthogonal principal modes. Unlike neural-network-based approaches, 3DVA assumes a Gaussian latent distribution and models heterogeneity within a linear framework. We used the 3DVA implementation in cryoSPARC (version 4.5). The analysis was performed with 3 variability components and a filter resolution of 6 Å. The resulting variability coefficients were used as latent coordinates for downstream analysis.

#### RECOVAR

RECOVAR [7] estimates a regularized covariance matrix of the heterogeneous 3D density distribution and performs kernel regression in the resulting principal component space to reconstruct conformational variability. We used the official implementation with default parameters. The model was applied with 10 principal components for latent embedding on each dataset.

#### OPUS-DSD

OPUS-DSD [6] disentangles dynamics and compositional heterogeneity in cryo-EM data using a deep learning framework with separate latent spaces and decoders for large-scale dynamics and compositional changes. We used the official implementation with default hyperparameters. The model was trained for 25 epochs with a latent dimension of 12 on each dataset.

#### CryoDECO

CryoDECO [8] leverages foundation model priors to improve heterogeneous cryo-EM reconstruction, enabling deconstructing of extreme compositional and conformational heterogeneity. We used the official implementation with default hyperparameters. The model was trained for 25 epochs with a latent dimension of 128 on each dataset.

#### 3DFlex

3DFlex [5] models continuous conformational flexibility by learning a non-rigid deformation field that maps a canonical density to each individual particle conformation. We used the 3D Flex implementation in cryoSPARC (version 4.5) with 2 latent dimensions and default parameters for both mesh preparation and training.

#### WAVE + CryoSPARC analysis

Based on the clustering and filtering results obtained via WAVE, the particle subsets corresponding to each cluster were imported into CryoSPARC(version 4.5). For each subset, an initial homogeneous reconstruction was performed to generate a preliminary three-dimensional volume. Subsequently, these volumes were subjected to non-uniform refinement, utilizing cross-validation-based regularization and anisotropic spatial smoothing to achieve high-resolution 3D reconstructions for each identified static state.

### Molecular dynamics simulation of integrin *αvβ*8

#### Construction of the integrin αvβ8 ectodomain model

The cryo-EM structure of integrin *αvβ*8 in complex with the Fabs C6D4 and 11D12v2 (PDB ID: 6UJB) was used as the initial experimental model. In the original study, an additional Fab, 11D12v2, was introduced to facilitate high-resolution reconstruction by binding to the *α*V thigh region. The authors noted that this Fab was used primarily to improve particle alignment and did not affect L-TGF-*β* binding or the conformation of the integrin headpiece [18]. Thus, 11D12v2 was not considered a functional component relevant to the conformational dynamics investigated here. In addition, the deposited coordinates for 6UJB contain only four chains that could be directly used for subsequent modelling, corresponding to integrin *αv*, integrin *β*8, the Fab C6D4 heavy chain, and the Fab C6D4 light chain, whereas no usable atomic coordinates were provided for 11D12v2. On the basis of both its auxiliary role in structure determination and the absence of a corresponding deposited atomic model, all subsequent structure reconstruction and molecular dynamics simulations were performed using only these four chains.

In 6UJB, the leg of chain A (integrin *αv*) and chain B (integrin *β*8) are incomplete. To obtain a complete ectodomain structure, we reconstructed the *αv* and *β*8 chains using AlphaFold 3, with a previously determined integrin *αvβ*3 structure (PDB ID: 8XEN) used as a structural template to guide modelling. The rebuilt *αv* and *β*8 chains covered both the headpiece and leg regions used in subsequent analyses and were then combined with the experimentally resolved Fab C6D4 chains from 6UJB to generate a complete integrin *αvβ*8-Fab ectodomain complex model. Because this study focused on the continuous extracellular conformational heterogeneity of the integrin leg, the transmembrane and intracellular regions were not included in the modelling or simulations.

#### System preparation

The reconstructed integrin *αvβ*8-Fab ectodomain complex was prepared using the Protein Preparation Wizard in Schrödinger Release 2025-4 [35]. During preparation, bond orders were assigned, hydrogen atoms were added, and the hydrogen-bonding network was optimized. The experimentally resolved Ca^2+^ and Mg^2+^ ions were retained throughout to preserve the native metal-coordination environment of the integrin ectodomain. Protonation states of hetero groups were assigned with Epik at pH 7.4 ± 2.0, followed by restrained minimization of the prepared structure.

The complex was solvated in TIP3P water in a rhombic dodecahedral box with a minimum solute-box distance of 1.2 nm, corresponding to a minimum separation of greater than 2.4 nm between the protein and its nearest periodic image. To mimic physiological conditions, the system was neutralized with counterions and supplemented with 0.15 M NaCl.

#### All-atom molecular dynamics simulations

All-atom molecular dynamics simulations were performed using GROMACS 2025.1 with the CHARMM36m force field [36, 37]. Following solvation and ion addition, each system was subjected to a two-step energy minimization procedure consisting of steepest-descent and conjugate-gradient minimization. After energy minimization, the system was equilibrated in two successive stages with positional restraints applied to the protein. First, a 1 ns NVT equilibration was carried out at 310 K with a 2 fs integration time step. Temperature was controlled using the V-rescale thermostat, with the protein and water-and-ions treated as separate coupling groups, each with a coupling constant of 1.0 ps. This was followed by a 1 ns NPT equilibration at 310 K and 1 bar using the same temperature-coupling scheme. Pressure was maintained using the C-rescale barostat with isotropic pressure coupling, a coupling time constant of 5.0 ps, and a compressibility of 4.5 × 10^-5^ bar^-1^. In both equilibration stages, non-bonded interactions were treated using the Verlet cutoff scheme; long-range electrostatics were calculated using the particle mesh Ewald (PME) method [38] with a real-space cutoff of 1.2 nm; and van der Waals interactions were treated using a force-switch scheme, beginning at 1.0 nm and reaching zero at 1.2 nm. Bonds involving hydrogen atoms were constrained using the LINCS algorithm [39].

Production simulations were subsequently carried out in the NPT ensemble after release of positional restraints, using the same temperature, pressure, and non-bonded interaction settings as in equilibration. To ensure extensive conformational sampling of the integrin *αvβ*8 leg, we initiated 20 independent trajectories from the same starting model using different random initial velocities. Each replicate was simulated for a total of 1 *µ*s, yielding an aggregate sampling time of 20 *µ*s. Together, these trajectories constituted the MD conformational ensemble used for subsequent landscape analysis. To exclude the initial relaxation phase, only the 200-1000 ns portion of each trajectory was used for quantitative analysis, with conformations sampled at 0.1 ns intervals. This yielded a total of 160,000 MD snapshots for subsequent landscape construction and quantitative comparison.

#### Convergence and coverage analysis of the MD conformational landscape

To evaluate the sampling exhaustiveness and convergence of the 20-*µ*s aggregate MD ensemble, we performed a structural saturation analysis. We quantified the conformational overlap between a subset of *k* independent trajectories (*k* ∈ [1, 19]) and the reference landscape defined by the complete 20-trajectory ensemble. For each value of *k*, *n* = 100 random combinations of trajectories were sampled to ensure statistical robustness. The conformational overlap (OVL) was calculated based on the similarity of the probability density distributions in the reduced dimensional space. The convergence was assessed by the mean overlap and its standard deviation (SD), where a near-unity overlap as *k* approaches 20 indicates that the ensemble has reached a state of sufficient sampling. Furthermore, we calculated the 95% confidence interval using the standard error of the mean (SEM) to confirm the stability of the observed convergence trend.

To evaluate the extent to which the MD-sampled ensemble encompasses the experimental cryo-EM conformational space, we performed a highest-density region (HDR) overlap analysis. Continuous probability density functions for both the MD trajectories and the cryo-EM-derived conformations (*n* = 10, 000 samples) were estimated using Gaussian kernel density estimation (KDE). The HDR for each ensemble was defined as the smallest region in the conformational space containing a given fraction of the total probability mass. We then quantified the inclusion of cryo-EM conformations within the MD landscape by calculating the fraction of the EM-defined HDR that was covered by the corresponding MD HDR across a range of probability mass thresholds (1% to 99%). This coverage was visualized as a function of the HDR threshold on a log_10_ scale to assess the representativeness of the MD ensemble across both highly populated states and surrounding structural fluctuations.

## 8 Data availability

The cryo-EM single particle datasets can be accessed on EMPIAR: integrin *αvβ*8 (EMPIAR-10345), KCTD5/CUL3^NTD^/G*βγ* (EMPIAR-11734), LIS1/dynein (EMPIAR-12715). The simulated benchmark datasets used in this work are available through CryoBench: Ribosembly, Tomotwin-100, and IgG-1D [19]. The pretrained weights of CryoUNI are available at https://doi.org/10.5281/zenodo.19513130. Results for the CryoBench dataset (IgG-1D, Ribosembly, and Tomotwin-100) are deposited at https://doi.org/10.5281/zenodo.19592117. Experimental results are available via https://doi.org/10.5281/zenodo.19592853 (EMPIAR-11734; KCTD5/CUL3*^NT^ ^D^*/G*βγ*), https://doi.org/10.5281/zenodo.19592956 (EMPIAR-12715;

LIS1-mediated Dynein), and https://doi.org/10.5281/zenodo.19592995 (EMPIAR-10345; Integrin *α*v*β*8).

## 9 Code availability

Source code of CryoUNI, pre-trained model weights, and training, inference scripts are available at https://github.com/Cellverse/cryouni.

## 10 Acknowledgements

We thank Y.-B. Shan and T. Hua for helpful discussions, and Z.-W. Luo and Y. Yan for guidance on using their baseline implementation. This work was supported in part by the National Natural Science Foundation of China (Grant No. W2431046), the Central Guided Local Science and Technology Foundation of China (Grant No. YDZX20253100001001), the National Key R&D Program of China (Grant No. 2025YFA1309603), the 2024 Shanghai Action for Science, Technology and Innovation Program of the Natural Science Foundation of Shanghai (Grant No. 24JS2820100 to Z.-J.L.), the MoE Key Laboratory of Intelligent Perception and Human-Machine Collaboration (ShanghaiTech University), the Shanghai Frontiers Science Center of Human-centered Artificial Intelligence, and the Shanghai Frontiers Science Center for Biomacromolecules and Precision Medicine at ShanghaiTech University. We acknowledge the High-Performance Computing Platform and the Bio-Electron Microscopy Facility of ShanghaiTech University for computational and infrastructure support.

---

**Extended Data Fig. 1.**
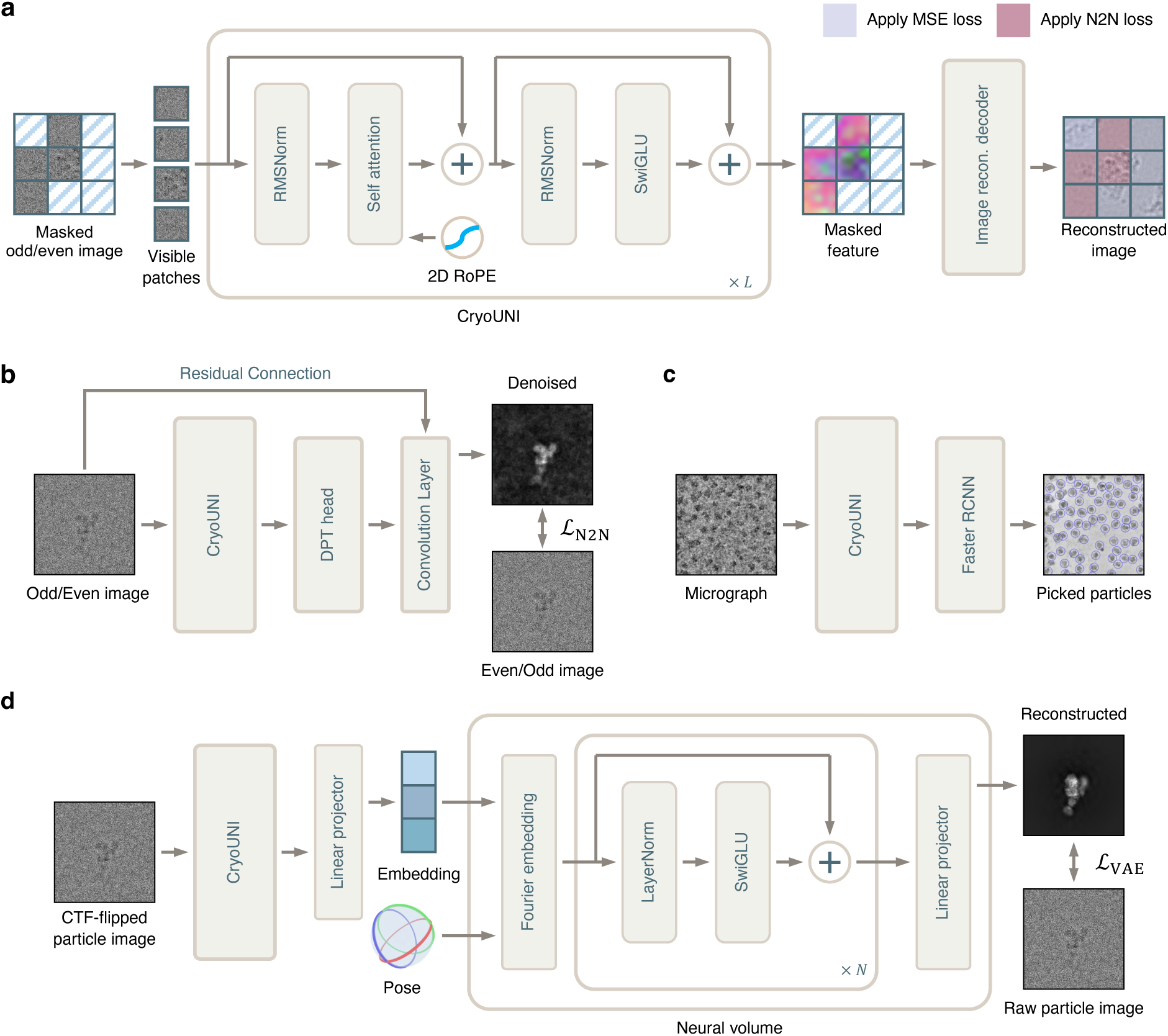
Network architecture of CryoUNI and its downstream applications. **a**, CryoUNI adopts a modern ViT architecture and undergoes a first-stage denoising-reconstruction pretraining based on the N2N and MAE frameworks. **b**, In the second-stage denoising pretraining, the model incorporates a DPT head while focusing on a specialized frequency-domain protection mechanism to preserve structural integrity. **c**, The particle picking model, built upon the Fast R-CNN object detection framework, is utilized in an iterative “picking-curating-retraining” cycle to generate the large-scale particle dataset. **d**, A CryoDRGN-style heterogeneous reconstruction framework with a specifically designed implicit neural representation volume decoder, which predicts slice intensities in the frequency domain from 3D coordinates and latent variables. Example images are from EMPIAR-10002 [42], EMPIAR-10076 [40], and EMPIAR-10345 [18].

**Extended Data Fig. 2.**
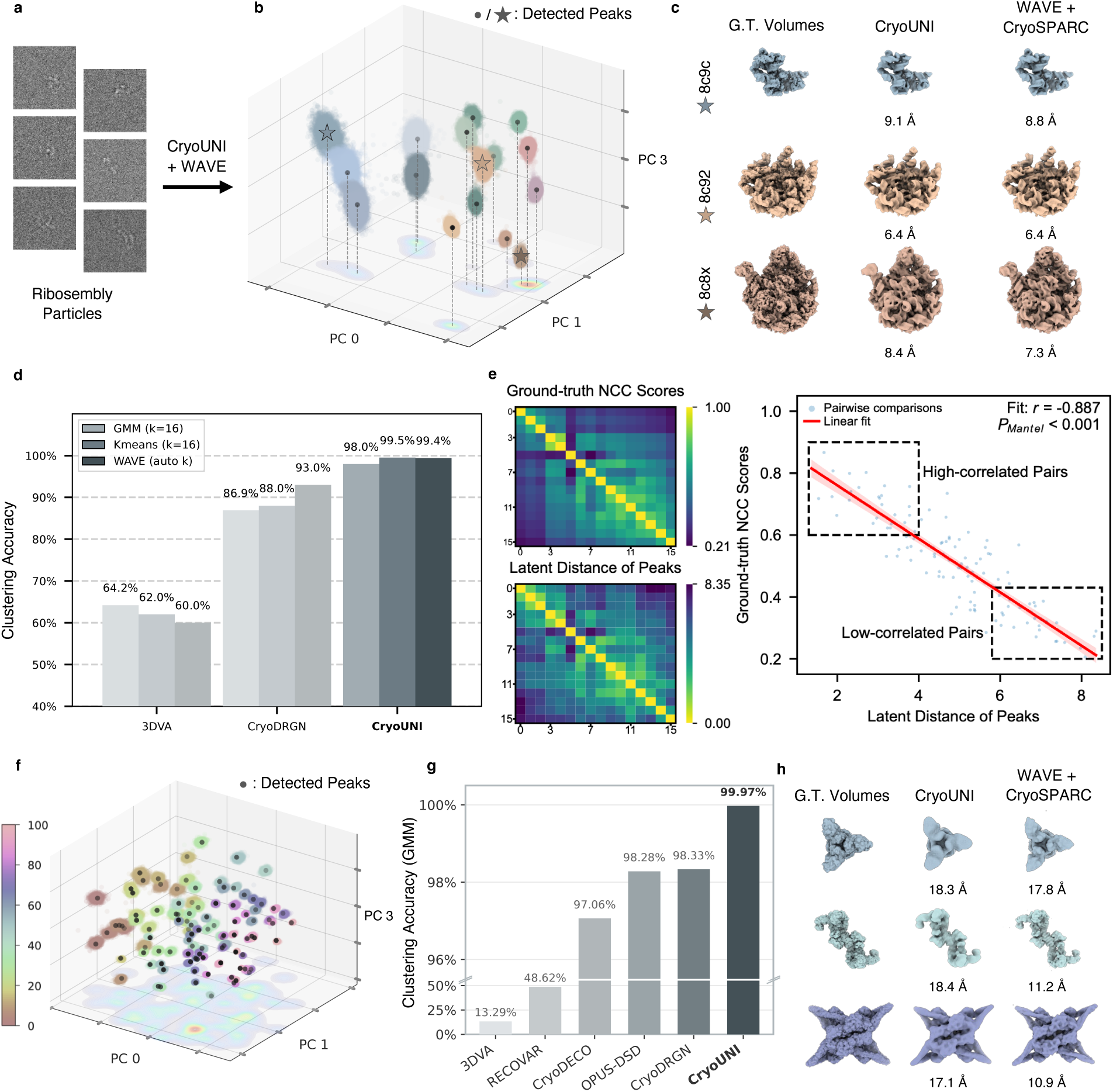
Evaluation of the latent landscape on compositional heterogeneity benchmarks. **a**, Example particle images from the Ribosembly dataset. **b**, Three-dimensional visualization of the CryoUNI latent landscape for the Ribosembly dataset (16 compositional states). Each high-density region corresponds to a distinct assembly intermediate. Detected peaks are marked by circles and stars. **c**, Representative reconstructed volumes for selected Ribosembly states (8c9c, 8c92, 8c8x). Ground-truth (G.T.) volumes are compared with CryoUNI decoder outputs and WAVE-guided cryoSPARC reconstructions, with achieved resolution indicated below each map. **d**, Clustering accuracy on the Ribosembly dataset across latent representations learned by 3DVA, CryoDRGN, and CryoUNI, using GMM (*k*=16), K-means (*k*=16), and WAVE (automatic *k*). **e**, Correspondence between latent distance and structural similarity. Top: ground-truth pairwise NCC matrix. Bottom: pairwise latent distance matrix between detected peaks. Right: scatter plot showing strong negative correlation between latent distance and structural similarity (Mantel test with 999 random permutations of the distance matrix; Pearson *r* = *−*0.887, *P*_Mantel_ *<* 0.001). **f**, Three-dimensional visualization of the CryoUNI latent landscape for the Tomotwin-100 dataset (100 structural classes), colored by class index. WAVE automatically detects all 100 density peaks. **g**, Clustering accuracy (GMM) on the Tomotwin-100 dataset across methods. **h**, Representative reconstructed volumes for selected Tomotwin-100 classes, comparing ground-truth volumes with CryoUNI and WAVE-guided cryoSPARC reconstructions.

**Extended Data Fig. 3.**
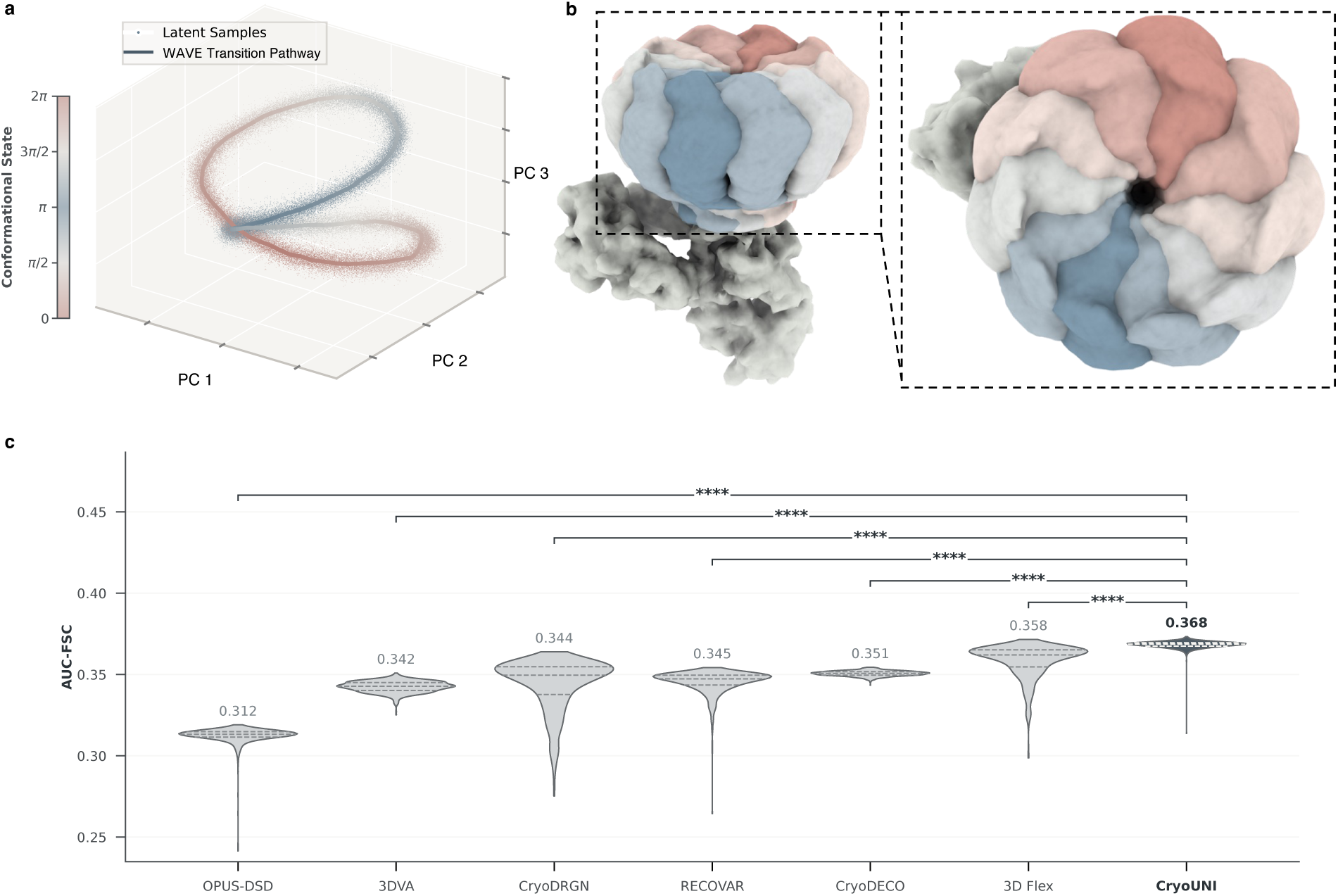
Evaluation of the latent landscape on the IgG-1D continuous heterogeneity benchmark. **a**, Three-dimensional visualization of the CryoUNI latent landscape for the IgG-1D dataset, colored by ground-truth conformational state (0 to 2*π*). The solid line denotes the trajectory traced by WAVE, recovering the full conformational cycle. **b**, Reconstructed volumes sampled along the WAVE trajectory, colored by conformational state as in **a**. Right: close-up view showing continuous structural variation across the conformational cycle. **c**, Per-conformation reconstruction consistency measured by AUC-FSC across methods (*n* = 1,000 reconstructed volumes per method, stratified as 10 per conformational class *×* 100 classes). Violin plots show the full distribution of AUC-FSC values; the white circle indicates the median, the thick bar spans the interquartile range (IQR, 25th–75th percentiles), and the thin line extends to 1.5*×* IQR. CryoUNI achieves the highest mean (0.368) with the smallest variance. Statistical significance was assessed using two-sided Mann–Whitney U tests (CryoUNI vs. each baseline: *P <* 10*^−^*^252^; see Methods for exact test statistics).

**Extended Data Fig. 4.**
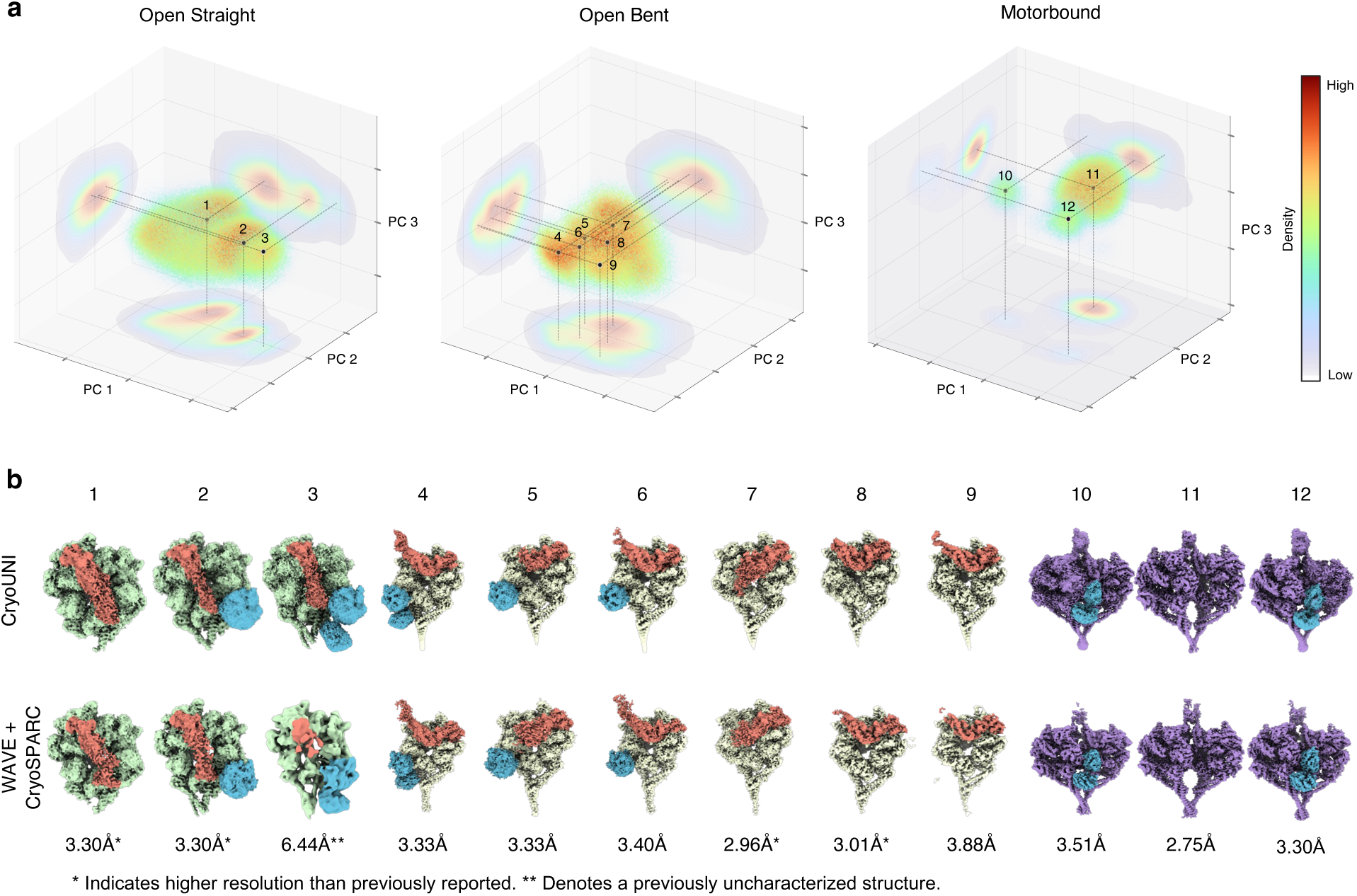
Hierarchical analysis of the conformational landscape for LIS1-mediated dynein activation. **a**, Local latent landscapes for the three major structural classes: open-straight (peaks 1–3), open-bent (peaks 4–9), and motor-bound (peaks 10–12). Multiple density peaks within each class indicate hierarchical sub-state organization. Dashed lines project peaks onto three PC planes; colored contours indicate relative energy thresholds (ΔΔ*G*r). **b**, Reconstructed volumes for all 12 identified sub-states. Top row: CryoUNI decoder outputs. Bottom row: WAVE-guided cryoSPARC reconstructions, with achieved resolution indicated below each map. Asterisks denote states with improved resolution compared to the original analysis (* moderate improvement; ** novel discovery).

**Extended Data Tab. 1.** Evaluation on CryoBench’s compositional datasets (Ribosembly and Tomotwin-100). Performance is measured by Adjusted Rand Index (ARI), Adjusted Mutual Information (AMI), and Accuracy (Acc.) on Ribosembly and Tomotwin-100 datasets. Upward arrows (*↑*) indicate higher values represent better performance. The best and second-best results across all methods are highlighted in boldface and underlined, respectively.

| Method | Clustering Algorithm | # Clusters | Ribosembly ($k = 16$) | | | Tomotwin-100 ($k = 100$) | | |
| --- | --- | --- | --- | --- | --- | --- | --- | --- |
| | | | ARI (↑) | AMI (↑) | Acc. ($\%, \uparrow$) | ARI (↑) | AMI (↑) | Acc. ($\%, \uparrow$) |
| RECOVAR | GMM | Pre-defined | 0.6751 | 0.8201 | 69.68 | 0.3383 | 0.6828 | 48.62 |
|  | K-means | Pre-defined | 0.7317 | 0.8512 | 74.89 | 0.3488 | 0.6592 | 48.58 |
|  | WAVE | Auto-detected | 0.7643 | 0.8945 | 77.67 | 0.0072 | 0.2884 | 13.77 |
| OPUS-DSD | GMM | Pre-defined | 0.8300 | 0.9199 | 78.18 | 0.9771 | 0.9923 | 98.28 |
|  | K-means | Pre-defined | 0.8656 | 0.9252 | 85.22 | 0.9776 | 0.9901 | 98.00 |
|  | WAVE | Auto-detected | 0.8032 | 0.9125 | 82.45 | 0.5626 | 0.8727 | 67.53 |
| CryoDRGN | GMM | Pre-defined | 0.8685 | 0.9021 | 86.88 | 0.9684 | 0.9877 | 98.33 |
|  | K-means | Pre-defined | 0.8730 | 0.9217 | 88.01 | 0.9702 | 0.9890 | 98.32 |
|  | WAVE | Auto-detected | 0.9072 | 0.9406 | 92.98 | 0.8506 | 0.9701 | 89.40 |
| 3DVA | GMM | Pre-defined | 0.5664 | 0.7221 | 64.18 | 0.0567 | 0.3325 | 13.29 |
|  | K-means | Pre-defined | 0.5460 | 0.7150 | 61.96 | 0.0575 | 0.3317 | 13.47 |
|  | WAVE | Auto-detected | 0.5652 | 0.7706 | 60.00 | 0.0015 | 0.0723 | 2.85 |
| CryoDECO | GMM | Pre-defined | 0.7082 | 0.8408 | 66.98 | 0.9751 | 0.9954 | 97.06 |
|  | K-means | Pre-defined | 0.8889 | 0.9277 | 86.57 | 0.9874 | 0.9977 | 98.57 |
|  | WAVE | Auto-detected | 0.7779 | 0.9081 | 78.42 | 0.1523 | 0.8357 | 57.90 |
| **CryoUNI** | GMM | Pre-defined | 0.9863 | 0.9812 | 97.97 | <u>0.9995</u> | <u>0.9996</u> | <u>99.97</u> |
|  | K-means | Pre-defined | **0.9919** | **0.9882** | **99.50** | **0.9995** | **0.9996** | **99.98** |
|  | WAVE | Auto-detected | <u>0.9907</u> | <u>0.9870</u> | <u>99.42</u> | 0.9992 | 0.9993 | 99.96 |

## Appendix A Pseudo-code of CryoUNI Encoding Process

Supplementary Alg. 1 shows the encoding process of CryoUNI from an input particle to its image feature.

### Algorithm 1

Image encoding process of CryoUNI

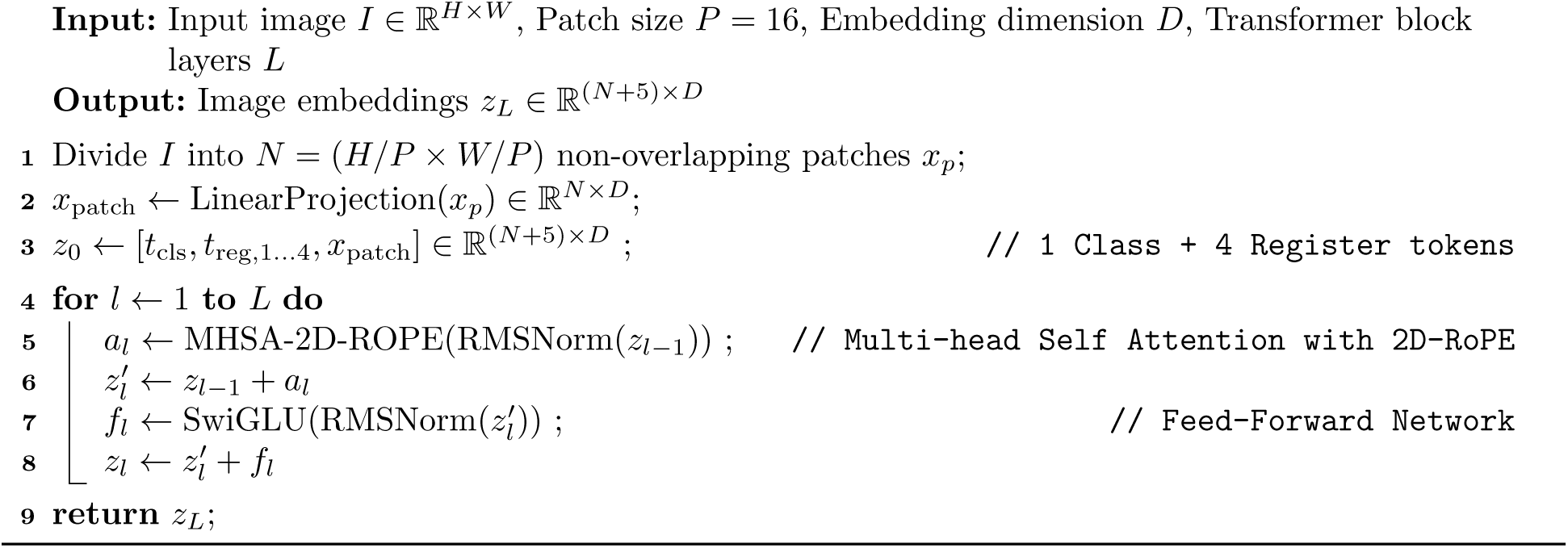

## Appendix B Qualitative Assessment of Pretraining

To evaluate the representational capacity of CryoUNI after pretraining, we visualize its performance across four single-particle datasets: EMPIAR-10023 [43], 10073 [44], 10345 [18], and 10980 [45]. As demonstrated in Supplementary Fig. C1, the model exhibits a robust ability to extract structural signal from particle images characterized by extremely low signal-to-noise ratios (SNR).

The latent semantics, captured by projecting the final-layer patch tokens onto their first three principal components, reveal a clear unsupervised segmentation between the molecular density and the surrounding noisy background. This behavior follows the feature visualization protocol proposed by Oquab et al. [46]. This emergent semantic clarity is complemented by the Multi-Head Self-Attention (MHSA) maps of the [CLS] token attention, which indicate that the model’s receptive field consistently converges on the central protein complex. By effectively suppressing stochastic background artifacts and prioritizing localized structural features, CryoUNI demonstrates that its self-supervised objective successfully distills high-fidelity biological representations without requiring explicit manual supervision.

## Appendix C Particle Dataset Curation

In the field of cryo-EM single-particle analysis, standardized and high-quality annotated data remain relatively scarce, posing significant challenges for training downstream supervised models. To address this, we adopted the “data engine” concept proposed by the Segment Anything Model [47] and implemented a collaborative ‘model-in-the-loop” dataset annotation strategy. Specifically, we designed an iterative optimization workflow: First, we constructed an initial training set through manual annotation and specialized software tools to train preliminary models. Subsequently, these models were leveraged to assist in large-scale semi-automated data annotation. This process not only expanded the scale of the dataset but also increased structural and compositional diversity. Through multiple iterative cycles, both dataset quality and quantity were progressively improved, leading to a synergistic enhancement in model robustness.

For the particle picking task, building upon curated high-quality micrographs, we conducted six rounds of iterative annotation cycles. This involved integrating multiple labeling tools (such as blob picking, template picking, and 2D classification [2]) and incorporating predictions from existing models like Topaz [48]. This strateg yielded a comprehensive corpus of 746 datasets, comprising 22,631,283 labeled particles.

### C.1 Particle Picking Model Training and Inference Details

The particle picking task aims to precisely identify target protein particles in highly noisy cryo-EM images. To adapt the micrograph version of CryoUNI for this task, we performed supervised fine-tuning using the Detectron2 framework [49], following the protocol of DRACO [17]. During training stage, the optimizer used was AdamW (*β*_1_ = 0.9*, β*_2_ = 0.999) with a base learning rate of 10^−4^ and weight decay of 0.1. The model was trained for 100 epochs with a batch size of 256. All other parameters follow the Detectron2 default configurations. The default aspect ratio is set to 1.0 with a non-maximum suppression (NMS) threshold of 0.6. During inference stage, we resize the micrograph into 1024 × 1024 and directly inference on it. A user specified score threshold (typically 0.0001) is used for filter out those low quality particles. The source of these particle dataset are listed in Supplementary Table C1.

**Supplementary Fig. C1.**
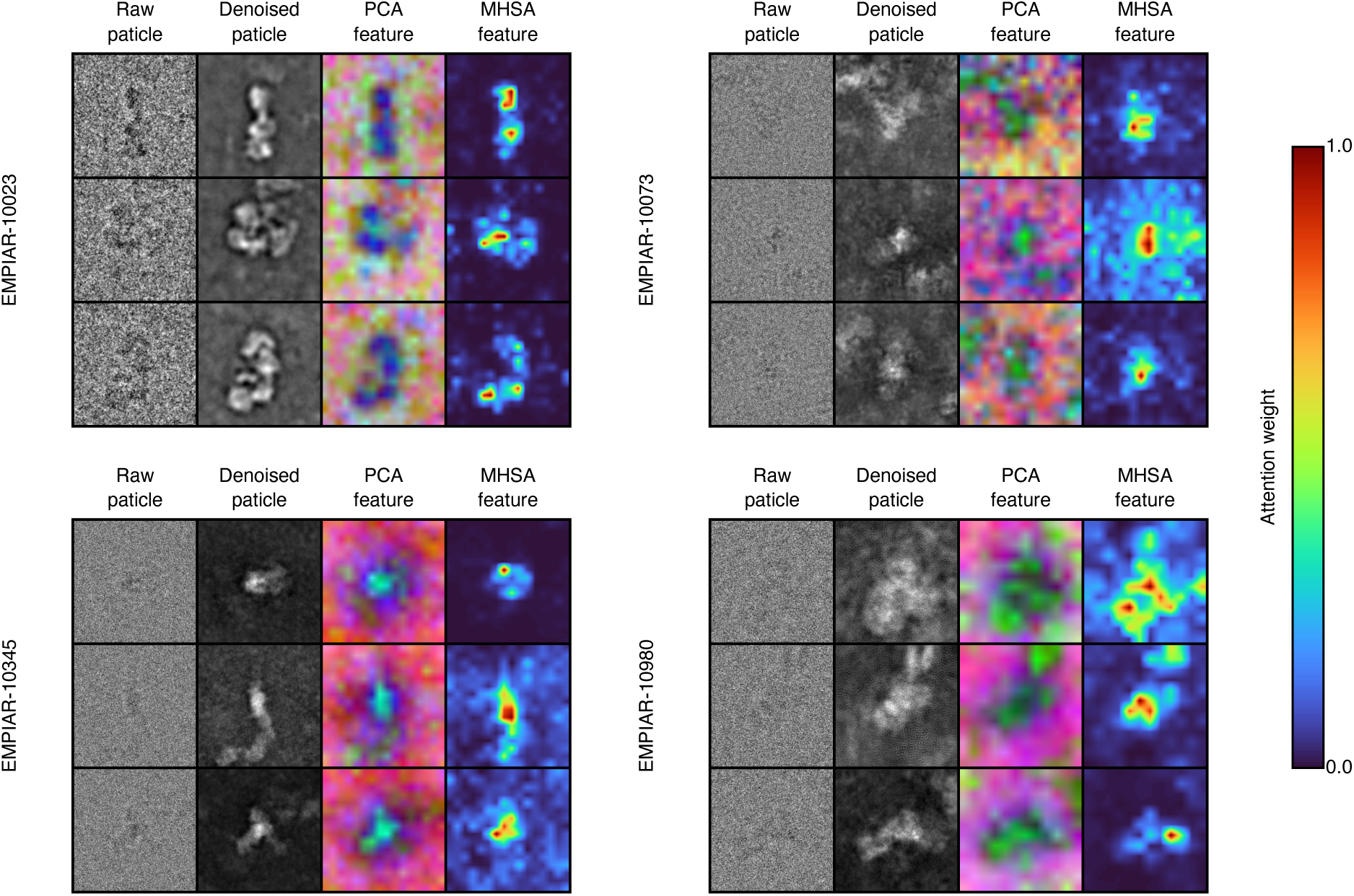
Visualization of Denoising and Latent Semantics. Qualitative results of CryoUNI across diverse Cryo-EM single-particle datasets (EMPIAR-10023 [43], 10073 [44], 10345 [18], and 10980 [45]). For each example, raw and denoised particles are shown alongside their patch token visualizations. The PCA features are generated by projecting the last layer’s patch tokens onto their first three principal components and mapping them to the RGB space (following the DINOv2 [46] protocol), where background signal is usually represented in red. The Multi-Head Self-Attention (MHSA) maps from the last layer illustrate the distribution of the model’s focus. Together, these visualizations demonstrate the ability of CryoUNI to distill meaningful structural signals from extremely noisy backgrounds, maintaining a consistent focus on target structures at the center of the image.

**Supplementary Fig. C2.**
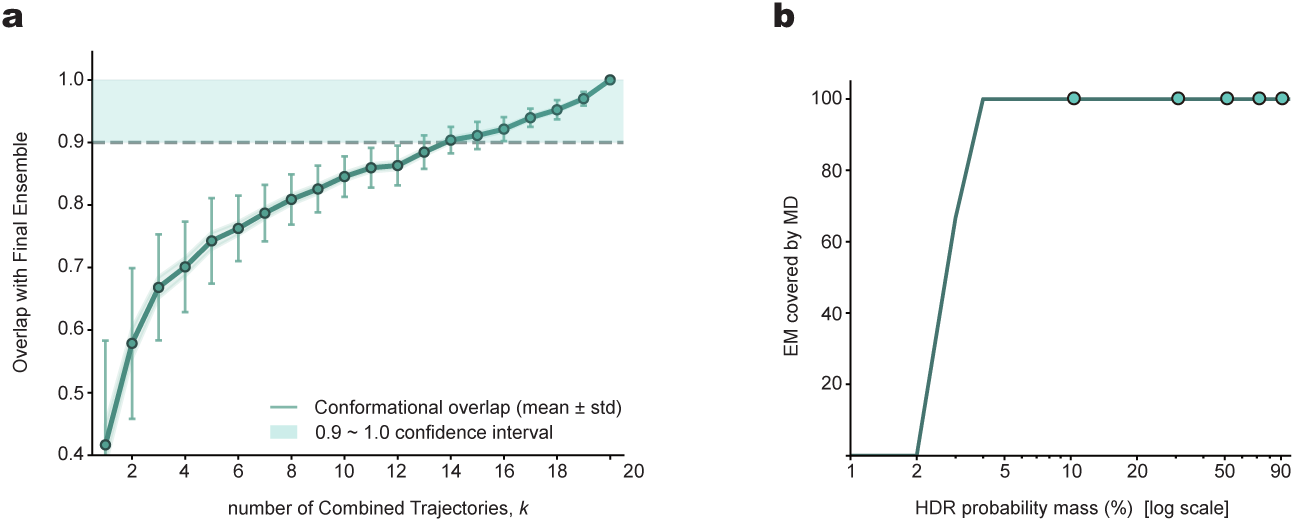
Convergence of MD simulations and coverage of CryoUNI conformations within the MD-derived landscape. - a, Saturation of the MD conformational ensemble. Overlap between landscapes from *k* random 1-*µ*s trajectories and the full 20-*µ*s ensemble. Error bars represent mean *±* s.d. computed from *n* = 100 random subsets for each value of *k*, drawn from the 20 independent trajectories. Near-unity overlap as *k →* 20 indicates convergence and sufficient sampling. b, Inclusion of CryoUNI conformations within the MD landscape. Fraction of CryoUNI samples (*n* = 10000) within the MD highest-density region (HDR) versus probability mass threshold (log_10_ scale). Nearly all CryoUNI conformations lie within the MD-sampled region across thresholds, confirming that cryo-EM captures the most populated states while MD explores surrounding fluctuations.

**Supplementary Tab. C1.** Pretraining dataset from EMPIAR.

| EMPIAR ID |  |  |  |  |  |  |  |  |  |  |  |
| --- | --- | --- | --- | --- | --- | --- | --- | --- | --- | --- | --- |
| 10002 | 10023 | 10058 | 10078 | 10084 | 10088 | 10117 | 10153 | 10154 | 10156 | 10181 | 10185 |
| 10186 | 10196 | 10200 | 10204 | 10206 | 10216 | 10218 | 10244 | 10247 | 10248 | 10249 | 10250 |
| 10251 | 10252 | 10257 | 10258 | 10259 | 10261 | 10263 | 10264 | 10265 | 10266 | 10269 | 10270 |
| 10275 | 10281 | 10284 | 10288 | 10292 | 10298 | 10303 | 10308 | 10309 | 10314 | 10317 | 10332 |
| 10334 | 10337 | 10338 | 10342 | 10351 | 10352 | 10359 | 10361 | 10366 | 10381 | 10382 | 10383 |
| 10384 | 10385 | 10386 | 10388 | 10390 | 10394 | 10400 | 10405 | 10408 | 10409 | 10410 | 10422 |
| 10423 | 10424 | 10428 | 10430 | 10431 | 10437 | 10438 | 10439 | 10440 | 10441 | 10444 | 10446 |
| 10456 | 10465 | 10469 | 10471 | 10474 | 10477 | 10480 | 10485 | 10486 | 10487 | 10488 | 10492 |
| 10496 | 10500 | 10501 | 10502 | 10503 | 10506 | 10507 | 10508 | 10509 | 10510 | 10511 | 10513 |
| 10518 | 10521 | 10522 | 10523 | 10524 | 10525 | 10526 | 10527 | 10529 | 10530 | 10535 | 10536 |
| 10540 | 10541 | 10543 | 10544 | 10546 | 10547 | 10549 | 10550 | 10552 | 10556 | 10557 | 10558 |
| 10559 | 10561 | 10565 | 10566 | 10568 | 10569 | 10571 | 10572 | 10573 | 10574 | 10575 | 10576 |
| 10577 | 10578 | 10579 | 10580 | 10581 | 10582 | 10583 | 10584 | 10585 | 10586 | 10588 | 10591 |
| 10594 | 10595 | 10596 | 10597 | 10598 | 10599 | 10600 | 10605 | 10607 | 10608 | 10609 | 10610 |
| 10611 | 10612 | 10626 | 10627 | 10628 | 10629 | 10635 | 10637 | 10639 | 10641 | 10642 | 10644 |
| 10645 | 10646 | 10647 | 10648 | 10649 | 10650 | 10658 | 10659 | 10660 | 10662 | 10665 | 10666 |
| 10671 | 10673 | 10674 | 10676 | 10680 | 10682 | 10683 | 10684 | 10691 | 10692 | 10695 | 10697 |
| 10698 | 10699 | 10708 | 10709 | 10711 | 10712 | 10714 | 10715 | 10716 | 10717 | 10721 | 10723 |
| 10725 | 10726 | 10728 | 10732 | 10733 | 10736 | 10737 | 10739 | 10741 | 10742 | 10743 | 10744 |
| 10745 | 10746 | 10747 | 10748 | 10749 | 10750 | 10754 | 10757 | 10758 | 10759 | 10763 | 10764 |
| 10774 | 10775 | 10776 | 10777 | 10780 | 10793 | 10795 | 10802 | 10803 | 10807 | 10808 | 10809 |
| 10810 | 10811 | 10813 | 10816 | 10817 | 10821 | 10823 | 10824 | 10826 | 10827 | 10828 | 10833 |
| 10834 | 10835 | 10836 | 10837 | 10838 | 10842 | 10843 | 10844 | 10845 | 10846 | 10848 | 10849 |
| 10850 | 10851 | 10852 | 10853 | 10854 | 10855 | 10856 | 10857 | 10858 | 10861 | 10863 | 10864 |
| 10865 | 10866 | 10867 | 10872 | 10875 | 10877 | 10878 | 10879 | 10880 | 10881 | 10886 | 10887 |
| 10892 | 10894 | 10896 | 10897 | 10899 | 10900 | 10901 | 10902 | 10904 | 10905 | 10909 | 10910 |
| 10911 | 10912 | 10913 | 10914 | 10915 | 10921 | 10924 | 10926 | 10927 | 10928 | 10931 | 10932 |
| 10936 | 10939 | 10941 | 10942 | 10945 | 10948 | 10949 | 10950 | 10951 | 10952 | 10953 | 10954 |
| 10955 | 10961 | 10964 | 10965 | 10966 | 10974 | 10975 | 10976 | 10977 | 10978 | 10979 | 10980 |
| 10984 | 10990 | 10991 | 10995 | 10996 | 10997 | 10999 | 11000 | 11006 | 11007 | 11008 | 11009 |
| 11010 | 11012 | 11013 | 11014 | 11021 | 11022 | 11029 | 11030 | 11031 | 11032 | 11045 | 11046 |
| 11047 | 11048 | 11049 | 11050 | 11051 | 11052 | 11053 | 11054 | 11056 | 11057 | 11059 | 11061 |
| 11062 | 11067 | 11068 | 11069 | 11071 | 11072 | 11073 | 11074 | 11075 | 11076 | 11079 | 11080 |
| 11081 | 11084 | 11091 | 11096 | 11097 | 11098 | 11099 | 11100 | 11101 | 11102 | 11105 | 11106 |
| 11107 | 11108 | 11109 | 11110 | 11117 | 11118 | 11119 | 11121 | 11124 | 11127 | 11132 | 11134 |
| 11135 | 11136 | 11137 | 11138 | 11152 | 11168 | 11170 | 11171 | 11176 | 11177 | 11178 | 11179 |
| 11180 | 11182 | 11186 | 11187 | 11189 | 11192 | 11195 | 11196 | 11197 | 11199 | 11202 | 11203 |
| 11205 | 11207 | 11208 | 11209 | 11210 | 11211 | 11212 | 11213 | 11222 | 11223 | 11224 | 11225 |
| 11229 | 11230 | 11231 | 11232 | 11234 | 11235 | 11236 | 11244 | 11245 | 11246 | 11247 | 11248 |
| 11249 | 11250 | 11251 | 11255 | 11256 | 11261 | 11265 | 11266 | 11272 | 11276 | 11279 | 11280 |
| 11281 | 11283 | 11285 | 11286 | 11287 | 11288 | 11289 | 11290 | 11291 | 11292 | 11294 | 11295 |
| 11296 | 11297 | 11298 | 11299 | 11304 | 11305 | 11307 | 11308 | 11309 | 11310 | 11311 | 11312 |
| 11313 | 11315 | 11316 | 11317 | 11321 | 11323 | 11326 | 11327 | 11328 | 11329 | 11330 | 11331 |
| 11332 | 11333 | 11334 | 11335 | 11336 | 11337 | 11338 | 11339 | 11340 | 11343 | 11345 | 11346 |
| 11347 | 11348 | 11349 | 11350 | 11351 | 11352 | 11353 | 11355 | 11356 | 11358 | 11359 | 11360 |
| 11361 | 11363 | 11364 | 11365 | 11371 | 11372 | 11373 | 11375 | 11377 | 11378 | 11379 | 11382 |
| 11385 | 11386 | 11387 | 11388 | 11389 | 11390 | 11391 | 11395 | 11396 | 11401 | 11402 | 11403 |
| 11409 | 11413 | 11422 | 11424 | 11427 | 11430 | 11431 | 11432 | 11433 | 11434 | 11435 | 11441 |
| 11448 | 11454 | 11455 | 11458 | 11459 | 11460 | 11461 | 11466 | 11471 | 11473 | 11474 | 11475 |
| 11476 | 11477 | 11481 | 11482 | 11483 | 11486 | 11487 | 11488 | 11489 | 11492 | 11493 | 11494 |
| 11495 | 11496 | 11498 | 11499 | 11506 | 11512 | 11513 | 11527 | 11530 | 11531 | 11532 | 11534 |
| 11535 | 11536 | 11539 | 11540 | 11541 | 11549 | 11551 | 11552 | 11558 | 11559 | 11560 | 11562 |
| 11563 | 11565 | 11566 | 11568 | 11570 | 11571 | 11572 | 11573 | 11574 | 11575 | 11583 | 11590 |
| 11592 | 11593 | 11597 | 11600 | 11601 | 11602 | 11606 | 11609 | 11611 | 11618 | 11624 | 11626 |
| 11628 | 11629 | 11630 | 11631 | 11632 | 11633 | 11634 | 11635 | 11640 | 11641 | 11642 | 11644 |
| 11645 | 11646 | 11648 | 11653 | 11654 | 11655 | 11656 | 11657 | 11659 | 11660 | 11661 | 11671 |
| 11672 | 11673 | 11675 | 11677 | 11682 | 11685 | 11689 | 11706 | 11712 | 11732 | 11733 | 11748 |
| 11749 | 11754 | 11762 | 11769 | 11770 | 11772 | 11773 | 11774 | 11775 | 11776 | 11784 | 11788 |
| 11790 | 11793 | 11799 | 11800 | 11807 | 11813 | 11814 | 11815 | 11823 | 11824 | 11825 | 11834 |
| 11836 | 11837 | 11838 | 11842 | 11846 | 11848 | 11852 | 11853 | 11855 | 11859 | 11866 | 11868 |
| 11869 | 11870 | 11871 | 11872 | 11878 | 11879 | 11886 | 11887 | 11890 | 11891 | 11893 | 11894 |
| 11912 | 11915 | 11916 | 11921 | 11926 | 11959 | 11980 | 11985 | 11994 | 12010 | 12023 | 12028 |
| 12029 | 12055 |  |  |  |  |  |  |  |  |  |  |

